# Non-invasive diagnosis of early chronic colitis cancerization via amplified sensing of miRNA-21 in NIR-IIb window

**DOI:** 10.1101/2024.12.19.629405

**Authors:** Fuheng Zhai, Baofeng Yun, Jiang Ming, Benhao Li, Tianyu Yu, Xiao Liu, Xusheng Wang, Zi-Han Chen, Changfeng Song, Mengyao Zhao, Jiyu Li, Aibin Liang, Fan Zhang

**Affiliations:** Department of Hematology, Shanghai Tongji Hospital, Tongji University School of Medicine, Shanghai 200065 (P. R. China); Department of Chemistry, State Key Laboratory of Molecular Engineering of Polymers, Shanghai Key Laboratory of Molecular Catalysis and Innovative Materials and iChem, Fudan University, Shanghai 200433 (P. R. China); Department of Oncology, Huadong Hospital, Fudan University, Shanghai 200433 (P. R. China); Department of Oncology, Pudong Hospital, Fudan University, Shanghai 200433 (P. R. China)

## Abstract

Early diagnosis of the colitis-associated colorectal cancer (CRC) is of great significance for improving prognosis and survival rates. However, the clinical used colonoscopy and biopsy methods are invasive and lack of sensitivity at early stage of cancerization. Herein, we present an amplified sensing strategy in the second near-infrared b (NIR-IIb, 1500-1700 nm) window for non-invasive in situ visualization of early cancerization biomarker miRNA-21. A CRC nanosensor composed of Er^3+^ doped lanthanide nanoparticles, DNAzyme, and IR820 dye is designed as a NIR-IIb ratiometric luminescence reporter, providing rapid feedback and high sensitive miRNA-21 detection with limit of detection (LOD) of 1.26 pM. This strategy enables a non-invasive detection of colitis-associated cancerization up to ∼4 weeks ahead of the clinically used biopsy analysis, providing a promising alternative for early diagnosis of chronic colitis cancerization and guidance for therapeutic management.

## Introduction

Colorectal cancer (CRC) is the second leading cause of cancer-related death in the world^1, 2^. Patients with inflammatory bowel disease are at increased risk of developing CRC^3-5^. In situ monitoring of the colitis-associated CRC transforming process is of great significance for elucidating the pathogenesis and behavior of inflammation-related cancers, which provides guidance for the early diagnosis of CRC^6, 7^. Colonoscopy is the currently most used method for diagnosing CRC and evaluating cancer progression^8^. However, this procedure is invasive with risk of perforation, bleeding, and other unpredicted complications. Moreover, the complex operation including requisite bowel preparation and the following biopsy of the polyps further exacerbates the time cost and economic burdens. Since most colitis-associated CRCs are slow growing and last for several years, the invasive colonoscopy and biopsy methods are not ideal approaches for the long-term monitoring of colitis-cancer transforming process. The growing understanding of the cellular physiological process has provided some candidate biomarkers as predictive tools for the early detection of cancers ^9, 10^. MicroRNA (miRNA) is a kind of small non-coding RNA molecules, composed of 19 to 25 nucleotides, which play an important role in the regulation of gene expression and immune-mediated disorders^11-13^. Several studies have shown that the cancerization of inflammatory bowel disease is associated with abnormal expression of miRNAs-21, which regulates biological behavior through the PTEN/PI-3 K/Akt signaling pathway in colorectal cancer cells^10, 13-15^ and is a promising biomarker in early diagnosis of CRC^12^. However, in vitro detection of CRC related miRNAs based on the fecal miRNA sequencing suffers from low specificity, and lack of spatial and temporal information^9^. Thus, accurate and real-time detection of miRNA in vivo is in urgent demand.

Luminescence imaging with fast feedback, non-ionizing radiation, and high sensitivity is revolutionizing both the fundamental biological research and clinical theranostic^16, 17^. In recent years, a number of strategies based on DNA biosensors for in vivo RNA luminescence imaging have been reported, enabling the in situ detection of RNAs in mice^18^. However, the spatiotemporal resolution of these methods is limited due to the significant effects of tissue scattering and absorption. In the past two decades, explosively developed the second near-infrared (NIR-II, 1000-1700 nm) window exhibits enhanced imaging contrast due to the decreased tissue absorption and scattering compared to the short-wavelength region (visible and NIR-I, 400-900 nm), facilitating the luminescence imaging from in vitro to in vivo. Additionally, the photon-bio-tissue interaction is negligible in the NIR-IIb (1500-1700 nm) subregion, offering a transparent window for imaging of physiological structures, and sensing of analytes including H^+^, adenosine triphosphate (ATP), enzyme, reactive oxygen species (ROS), and glutathione (GSH) under deep tissue penetration. Furthermore, using responsive NIR-II ratiometric luminescence, which introduces a self-calibrate signal during imaging, can circumvent the problems of signal fluctuation induced by probe concentration variation, thus enabling more accurate in vivo detection^19, 20^.

Herein, we report an amplified sensing strategy for noninvasively monitoring colitis-associated CRC transforming process in NIR-IIb window via a lanthanide-DNAzyme-dye hybrid ratiometric luminescence nanosensor (denoted as CRCsensor). The CRCsensor is constituted of three components: Er^3+^ doped lanthanide nanoparticle (ErNP) with NIR-IIb emission at 1550 nm under both 808 and 980 nm laser excitation (denoted as L_808_ and L_980_, respectively), DNAzyme locked with a miRNA-21 pairing short strand, and organic IR820 dye (λ_ex_=820 nm) conjugated DNAzyme substrate (sub-IR820) which competitively absorbed the 808 nm excitation laser and thus quenched the L_808_ signal of ErNP (Scheme 1A). Specifically, the locker stand (cyan chain) contains a miRNA-21 recognition segment and a DNAzyme complementary segment, which deactivates the DNAzyme. Meanwhile, the substrate is a DNA-RNA chimeric nucleic acid strand comprising a scissile adenosine ribonucleotide (rA) flanked by two DNA domains capable of binding to the DNAzyme. In the early stage of orthotopic CRC lesion, the overexpressed miRNA-21 competitively pairs with the locker DNA strand, peels off the locker strand, and activates the DNAzyme to cleave the adjacent sub-IR820 successively, thus releasing the quencher IR820, and finally amplifying the NIR-IIb L_808_ signal. During the miRNA-21 recognition, L_980_ signal remains consistent given that absorption of IR820 is negligible at 980 nm, providing a self-calibrated in the NIR-IIb ratiometric luminescence imaging to obtain the accurate in vivo miRNA-21 detection (Scheme 1B). Notably, the CRCsensor is introduced noninvasively from the anus to the colorectum, which is an effective and secure method compared to the conventional routes of intravenous or oral administration. With the LOD of 1.26 pM, the highly sensitive CRCsensor demonstrated the capacity to discern the presence of early-stage CRC leison in DSS-induced chronic colitis mice model, providing superior alternative for longterm monitoring and early diagnosis of the potential cancerization.

**Scheme 1.**
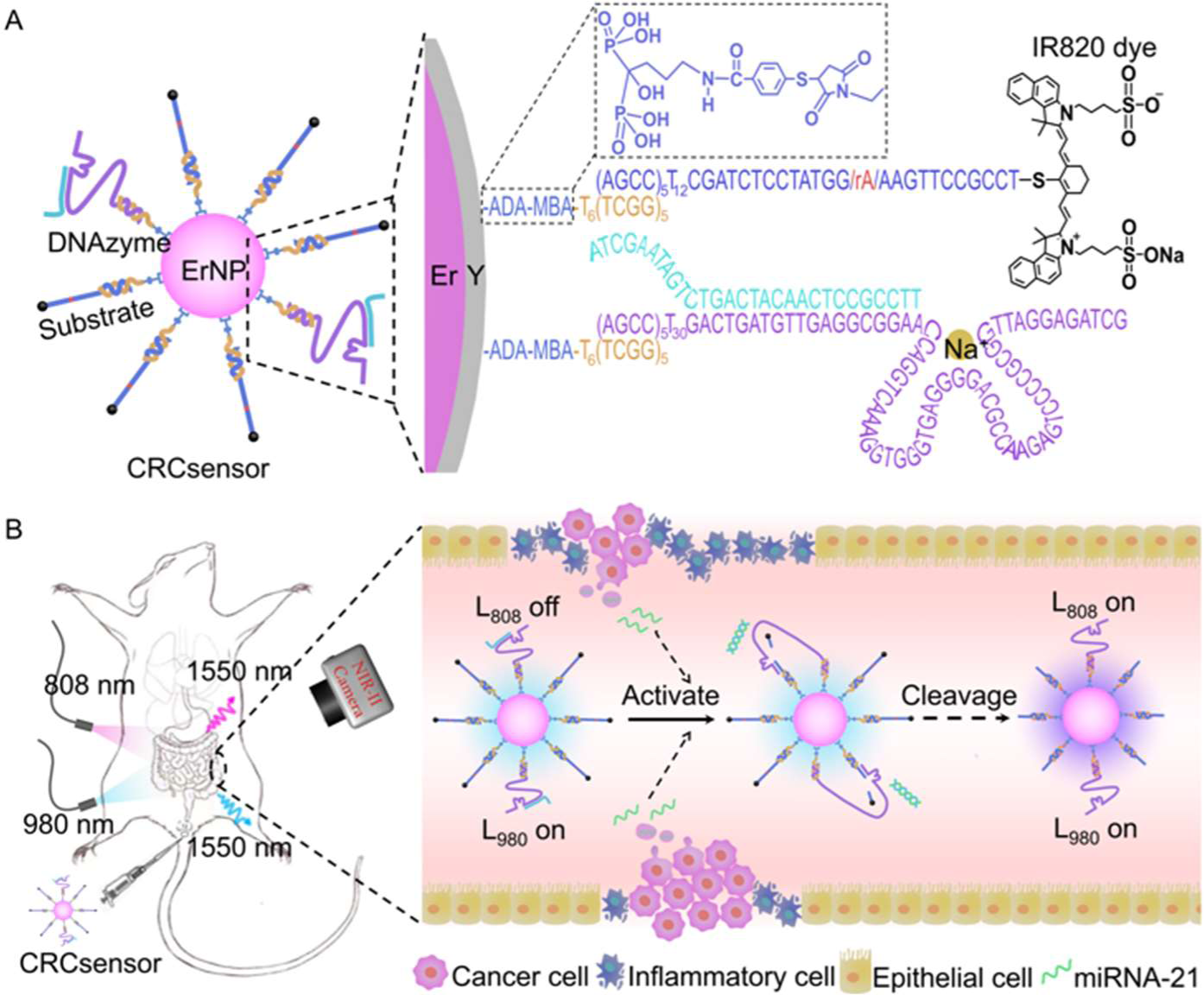
Schematic illustration of CRCsensor composition (A) and miRNA-21 sensing strategy in orthotopic CRC-bearing mice models (B).

## Results and Discussion

The synthesis flow of the CRCsensor was shown in Figure 1A. Hexagonal-phase NaErF_4_@NaYF_4_ nanoparticles (ErNPs) were synthesized through solvothermal methods. The diameter of the inner NaErF_4_ cores and the core-shell NaErF_4_@NaYF_4_ were 37.9 ± 0.7 nm and 62.9 ± 1.4 nm, respectively (Figure 1B, Figure S1-S4). The as-synthesized ErNPs showed distinct emission at 1550 nm under both 808 and 980 nm laser excitation (Figure S5). Subsequently, the alendronate sodium trihydrate (ADA) and 4-mercaptobenzoic acid (MBA) were covalently linked and modified on the surface of ErNPs via ligand exchange process. A maleimide-functionalized DNA strand (termed as linker DNA) was conjugated to the ErNP@ADA-MBA particles via thiol-Michael addition reaction, providing linkage sites for DNAzyme and sub-IR820. The successful modification was confirmed by the characteristic absorption peaks of ADA at 1120 cm^-1^ corresponding to stretching vibration of P=O and DNA characteristic absorption peak between 1600∼1700 cm^-1^ corresponding to stretching vibration of C=O and C=N of purine and pyrimidine in Fourier Transform Infrared (FTIR) Spectra (Figure 1D). The hydrodynamic size increasing from 63.3 ± 2.5 nm to 77.2 ± 4.4 nm, and zeta potential changing from 34.6 ± 5.7 mV to -30.0 ± 1.3 mV also validated the successful construction of the ErNP@ADA-MBA@linker DNA (Figure S6).

**Figure 1.**
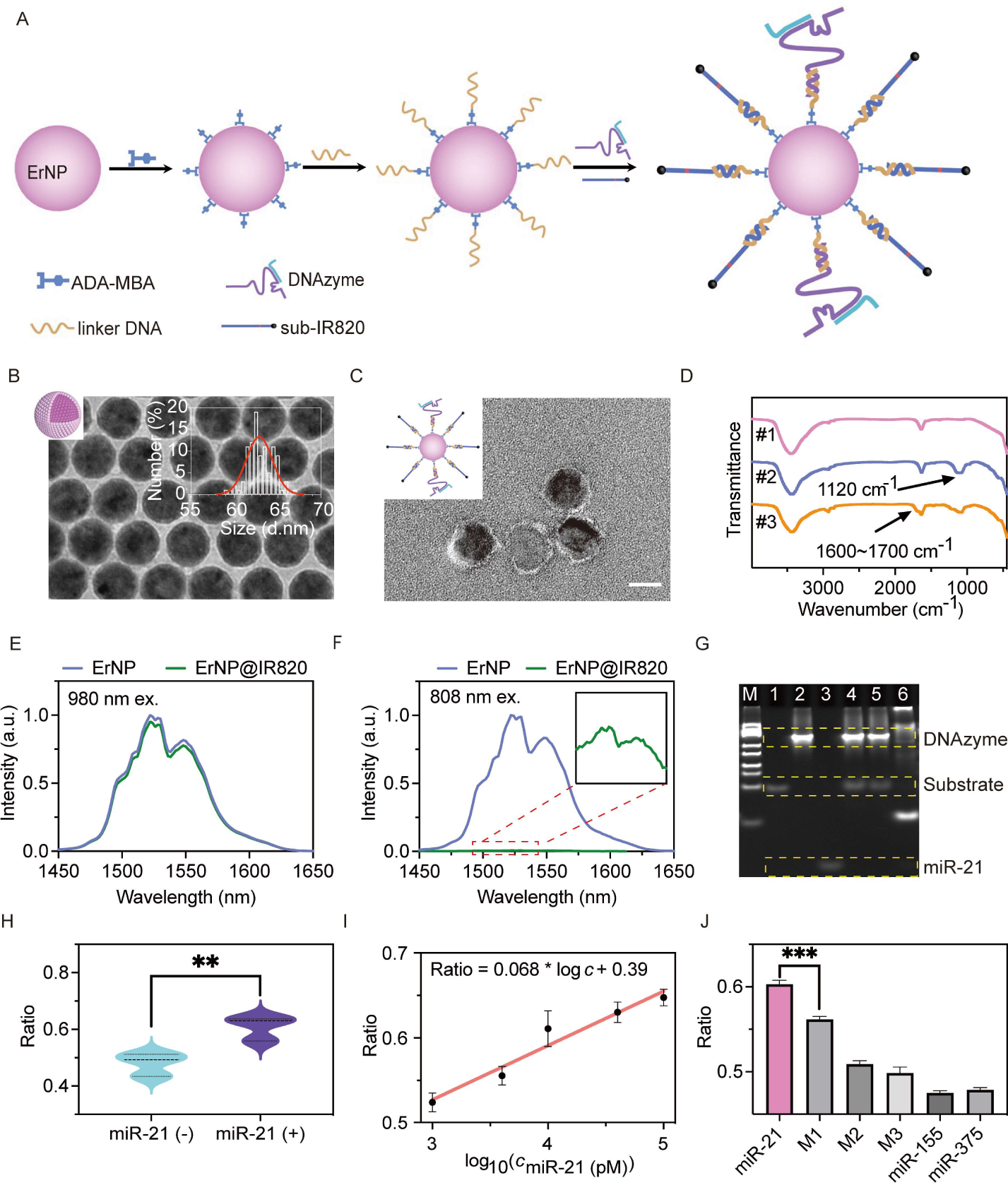
A) Schematic illustration of the CRCsensor fabrication. B) Transmission electron microscopy (TEM) image of the as-synthesized ErNP. Inset: size distribution, scale bar: 50 nm. C) TEM image of CRCsensor stained with 2% uranium acetate, scale bar: 50 nm. D) Fourier Transform Infrared (FTIR) Spectra of ErNP@BF_4_^-^ (#1), ErNP@ADA-MBA (#2), ErNP@ADA-MBA@linker DNA (#3). E, F) Luminescence intensity of the ErNP and the ErNP@IR820 under 980 nm (E) and 808 nm (F) excitation. G) Native polyacrylamide gel electrophoresis (PAGE) characterization of the DNAzyme activation and miRNA-21 recognition. The lanes in PAGE image are marker (lane M, 25 bp-500 bp), substrate (lane 1), DNAzyme (lane 2), miRNA-21 (lane 3), substrate + DNAzyme (lane 4), substrate + DNAzyme + Na^+^ (lane 5) and substrate + DNAzyme + Na^+^ + miRNA-21 (lane 6), respectively. H) L_808_/L_980_ ratio of the CRCsensor before and after incubation with miRNA-21. I) Calibration curves of the L_808_/L_980_ luminescence ratio of the CRCsensor in response to 1-100 nM miRNA-21. J) The L_808_/L_980_ luminescence ratio in response to miRNA-21, single mismatched miRNA-21 (M1), double mismatched miRNA-21 (M2), triple mismatched miRNA-21 (M3), miRNA-155 and miRNA-375, respectively. P values are indicated as follows: <0.05 (*), <0.01 (**), <0.001 (***), <0.0001 (****).

To achieve NIR-IIb sensing, the luminescence intensity at 1550 nm of the ErNPs under 808 nm excitation was regulated by the IR820 dye through the previously reported absorption competition-induced emission by our group^[10b]^ (ACIE, Figure S7). The absorption of the IR820 fluorophore at 808 nm was approximately ∼200000 times higher than that of the ErNPs attributed to the almost 5 orders of magnitude larger molar extinction coefficient of IR820 than ErNP (Figure S8). Given that IR820 exhibited no emission beyond 1500 nm, it can be employed as an effective photon-filtering layer of 808 nm excitation, causing significant luminescence quenching at 1550 nm when coated on the ErNP (Figure 1E, 1F, Figure S9). Thus, the luminescence of the CRCsensor at 1550 nm can only be excited by the 980 nm laser rather than the 808 nm laser. To obtain the sub-IR820, the IR820 dye was modified on the substrate via a substitution reaction involving the chlorine atom of IR820 and the sulfhydryl group of the 3’ terminus of the substrate, which was verified by the characteristic absorption peaks of the DNA strand at 260 nm and the IR820 dye at 820 nm in the UV spectra (Figure S10). Finally, sub-IR820 and DNAzyme were co-immobilized on the ErNPs via Watson-Crick base pairing principle, obtaining the well-dispersed CRCsensor with diameter size of 72.5 ± 2.2 nm (Figure 1B, Figure S11-S12). With 294 substrate and 94 DNAzyme on each CRCsensor, the luminescence ratio under 808- and 980- nm excitation channel markedly diminished 34% comparison to ErNP (Figure S13-14).

As the key sensing module, the miRNA-21 initiated DNAzyme-catalytic reaction were first investigated in aqueous solution by native polyacrylamide gel electrophoresis (PAGE, Figure 1G). In lane 6, in the present of miRNA-21, the corresponding band was not observed as that in lane 3, meanwhile, the disappeared substrate band further indicating that miRNA-21 lead to the hybridization and peeled off of the locker strand, activating the DNAzyme and the following catalytic reaction. Subsequently, the feasibility of the CRCsensor for the sensing of miRNA-21 in vitro was investigated (Figure 1H). Following the activation of the DNAzyme by miRNA-21 and the release of IR820 fluorophores from the ErNPs, the L808 of CRCsensor was restored, accompanied by a significant increase in the L808/L980 ratio value by approximately 16%. A gradual increase in L808/L980 ratio was observed in linear correlation with the concentration of miRNA-21 (1 nM-100 nM), providing a limit of detection (LOD) of 1.26 pM (Figure 1I), which was comparable to the previously works ^21, 22^. Meanwhile, to validate the sensing specificity of the CRCsensor, a series of nonspecific miRNAs, including single base mismatched miRNA-21 (M1), double base mismatched miRNA- 21 (M2), triple base mismatched miRNA-21 (M3), miRNA-155, and miRNA-375 were incubated with CRCsensor, respectively (Figure 1J). The results demonstrated that all miRNAs exhibited a minimal L808/L980 ratio increase in comparison to the target miRNA-21. Importantly, the mismatched M1 can be well distinguished from the target, demonstrating the high selectivity of CRCsensor. Furthermore, the significant increase in L808/L980 ratio after responding to miRNA-21 in saline, artificial colon fluid (ACF) and fetal bovine serum (FBS) illustrating the capability of sensitive and specific sensing in the complex microenvironment, providing the promising the in vivo application potential of CRCsensor (Figure S15).

Next, to investigate the sensing capability of CRCsensor towards endogenous miRNA, murine colon cancer cells (MC38, high miRNA-21 expressing) and immortalized murine colonic mucosal epithelial cells (MCEC, low miRNA-21 expressing) were selected as model cells (Figure 2A). First, the total miRNAs in MC38 and MCEC were extracted and incubated with CRCsensor, the 1.6-fold increase of L808/L980 ratio in MC38 lysates than MCEC lysates validated the successful responsiveness towards the endogenous miRNA (Figure 2B, 2C). Then, the CRCsensor was incubated with MC38 and MCEC cells for 4 h, followed by NIR-II luminescence microscope imaging with widefield mode. The acquired images of MC38 cells incubating with the CRC sensor showed an apparent recovery of luminescence in the 808 nm excitation channel, resulting in a 1.9-fold increase of L808/L980 ratio compared to the control MECE cells (Figure 2D, 2E, 2F). Furthermore, the MC38 cell and MCEC cell were treated with miRNA-21 mimic (DNA strand with same sequence as miRNA-21) and miRNA-21 inhibitor (DNA strand with complementary sequence as miRNA-21) respectively to up-regulate and down-regulate intracellular miRNA-21 expression to mimic miRNA different expression levels in cells. Compared to untreated cells, the miRNA-21 mimics treated MC38 and MCEC cells exhibited markedly stronger intracellular NIR-II luminescence (3.1- and 2.2- folds, respectively), whereas miRNA-21 inhibitor treated MC38 and MCEC cells displayed significantly reduced luminescence (0.54- and 0.71- fold, respectively), further validating the intracellular miRNA-21 responsive capacity of the CRCsensor (Figure S16-S17). The biocompatibility of the CRCsensor was also assessed by the CCK8 assay, which shows over 92% and 83% cell viability when the concentration of CRCsensor was below 50 μM (Figure S18).

**Figure 2.**
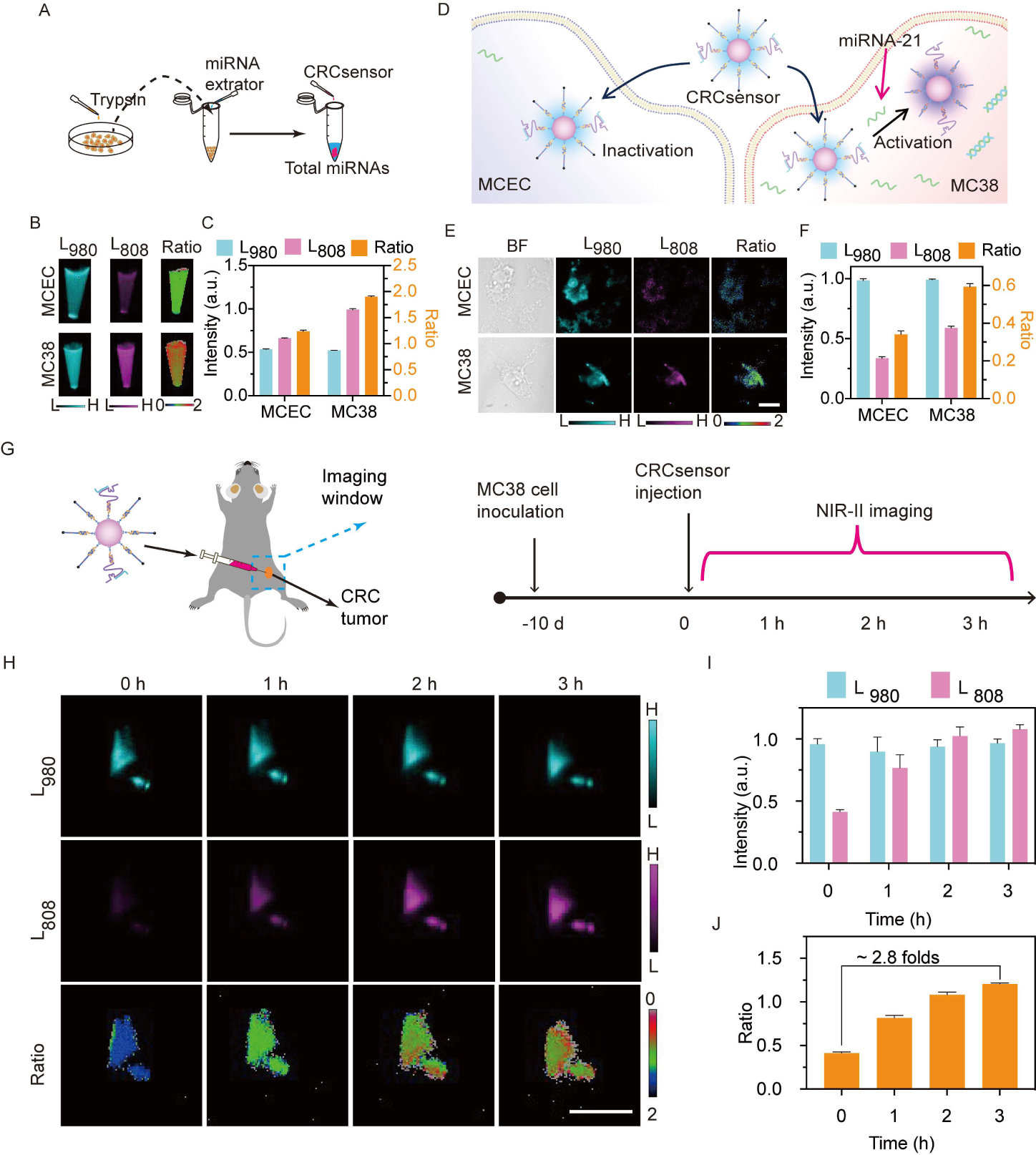
A) Schematic illustration of the total miRNA extraction and detection with CRCsensor. B) NIR-II luminescence and ratiometric images of the CRCsensor incubated with the total miRNAs extract from MCEC and MC38 cells, respectively (cell number: ∼10^7^). C) NIR-II luminescence intensity of CRCsensor from MCEC and MC38 cell lysates under 808- and 980- nm laser excitation and the corresponding calculated ratio values. D) Schematic illustration of MECE and MC38 cell incubated with CRCsensor. E) Representative NIR-II luminescence and ratiometric images of MCEC and MC38 cells incubated with the CRCsensor for 4 h. Scale bars: 25 μm. F) NIR-II luminescence intensity of CRCsensor from MCEC and MC38 cells under 808- and 980- nm laser excitation and the corresponding calculated ratio values. G) Schematic illustration of the NIR-II ratio luminescence imaging of subcutaneous CRC-bearing mice model. H) Representative in vivo NIR-II luminescence and ratiometric images of the subcutaneous CRC-bearing mice. Scale bars: 1.0 cm. I) NIR-II luminescence intensities of the CRCsensor from the subcutaneous CRC-bearing mice under 808- and 980- nm laser excitation. J) the calculated ratio values of the CRCsensor from the subcutaneous CRC-bearing mice under 808- and 980- nm laser excitation.

Subsequently, C57BL/6J mice bearing subcutaneous CRC tumors were selected as the model mice to demonstrate the application of the CRCsensor for in vivo miRNA-21 imaging (Figure 2G). As a control, a miNRA-21 non-responsive CRCsensor was designed and characterized (Figure S19-S20). A gradually luminescence recovery in the 808 nm channel was observed in the mice injected with the CRCsensor, while the L808 of the mice injected with the non-responsive CRCsensor exhibited minimal change after the injection (Figure 2H, 2I, Figure S21A-B). The CRCsensor injected group exhibited ∼2.8 folds increase of the L808/L980 ratio value while the control group showed slightly increase of L808/L980 ratio value (∼1.1 folds) at 3 hours after injection (Figure 2J, Figure S21C). These results indicated that the CRCsensor was capable of detecting miRNA-21 in vivo.

As the aberrant expression of miRNA-21 in the feces has been demonstrated to serve as an efficacious indicator for the CRC, we sought to ascertain the capacity of the CRCsensor for non-invasively detecting miRNA-21 in the colorectum of orthotopic CRC-bearing C57BL/6J mice (Figure 3A). Healthy C57BL/6J mice were used as control group. To facilitate the effective and secure delivery of the CRCsensors into the colorectum, an alternative approach was adopted whereby the CRCsensor was introduced noninvasively from the anus to the colorectum, as opposed to the conventional routes of intravenous or oral administration. As illustrated in the Figure 3B, upon administration of the CRCsensors into the mice colorectum, NIR-II luminescence signals of L980 and L808 were successfully collected in the abdomen of both CRC mice and healthy mice, demonstrating that the CRCsensor possess adequate tissue penetration and are suitable for miRNA-21 imaging in the colorectum tract in vivo. It should be noted that for in vivo imaging, intestinal peristalsis and alterations in intestinal position have the risk to yield inaccurate results when used for the monitoring of disease. In this study, following the delivery of the CRCsensor to the colorectum, the F808/F980 ratio luminescence in CRC-bearing mice at 4 hours yielded a self-calibrated result that was ∼1.6-fold higher than that observed at 0 hours, indicating that the CRCsensors exhibited a notable responsiveness to miRNA-21 in the colorectum tract (Figure 3C, 3D). Furthermore, the occurrence of CRC was confirmed by means of colorectal dissection images and a classic in vitro Hematoxylin-Eosin (H&E) analysis (Figure 3E). As for the healthy mice, no significant alteration was observed in the L808/L980 ratio over time (Figure 3G, 3H, 3I). No evidence of cancerization was observed in the colorectal dissection images and the H&E analysis of the healthy mice, which was consistent with the imaging results (Figure 3J). Subsequently, the mice were sacrificed and the colorectum tissues was imaged under 808 nm and 980 nm excitation lasers. The L808/L980 ratio of CRC-bearing mice colorectum tissue was ∼ 1.8 folds higher than that of the healthy mice (Figure S22). The in vivo ratiometric luminescence imaging results were in good agreement with the in vitro results, which can be attributed to the reduced scattering and deep penetration ability of CRCsensor NIR-IIb luminescence. The distribution of the CRCsensor in the organs of both the CRC-bearing mice and healthy mice demonstrated that the CRCsensors were exclusively present in the gastrointestinal tracts, indicating that they were not captured by organs and thus exhibited favorable safety characteristics (Figure 3F, 3K, Figure S23). It is noteworthy that the luminescence imaging strategy for non-invasive miRNA-21 detection in the colorectum is a more straightforward and expedient approach than the detection of miRNA-21 in feces in vitro.

**Figure 3.**
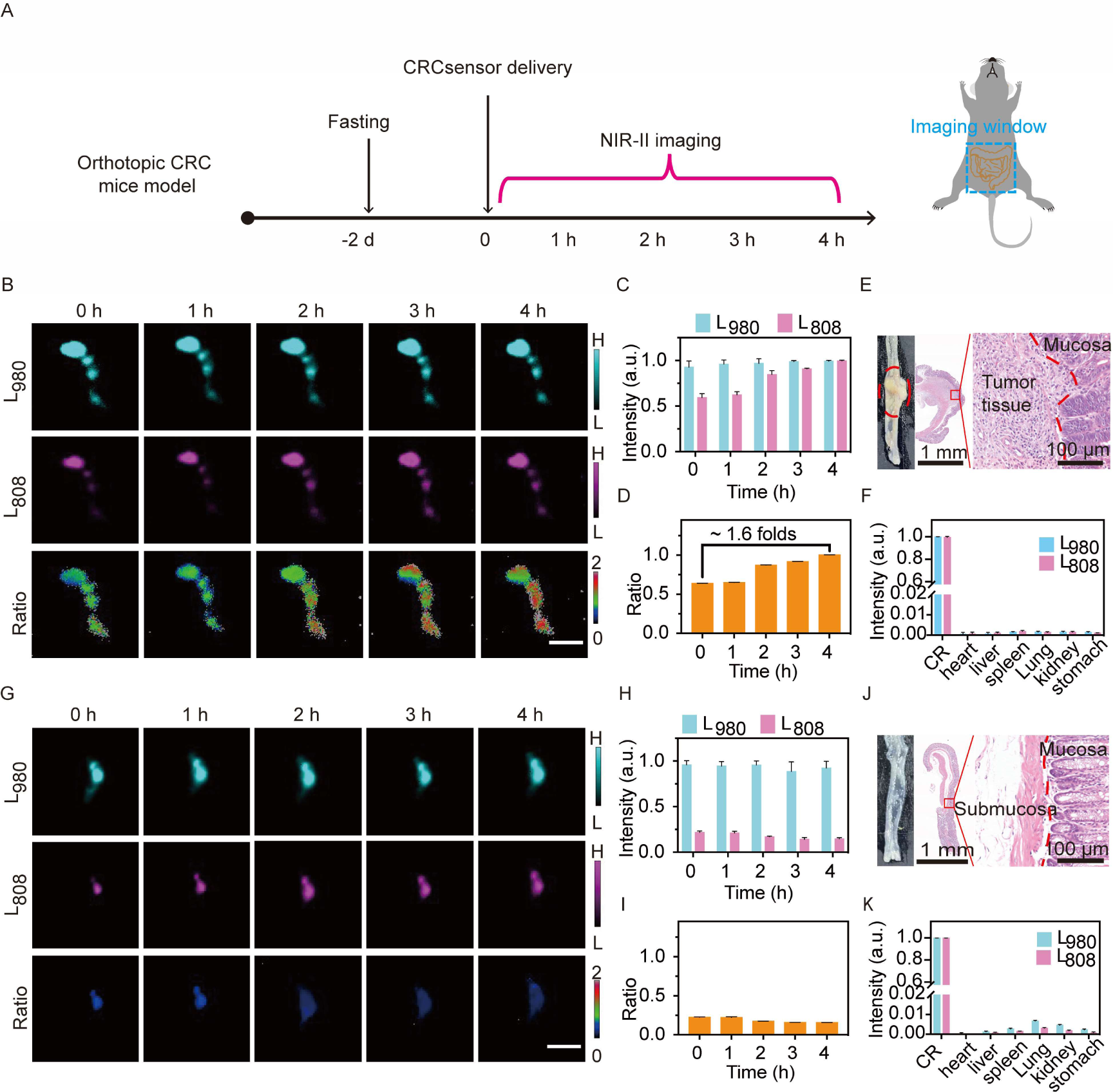
A) Schematic diagram of NIR-II ratiometric luminescence imaging of the orthotopic CRC mice model. B) Representative in vivo NIR-II luminescence and ratiometric images of the CRC-bearing mice. Scale bars: 1.0 cm. C-D) NIR-II luminescence intensity of the CRCsensor from CRC-bearing mice under 808- and 980- nm laser excitation (C) and the corresponding calculated ratio values (D). E) The colorectum dissection image (left) and Hematoxylin-Eosin (H&E) results (right) of the CRC-bearing mice. F) Luminescence intensity of the CRCsensor from different organs of the CRC-bearing mice under 808- and 980- nm laser excitation. G) Representative in vivo NIR-II luminescence and ratiometric images of the healthy mice. Scale bars: 1.0 cm. H-I) NIR-II luminescence intensity of the CRCsensor from healthy mice under 808- and 980- nm laser excitation (H) and the corresponding calculated ratio values (I). J) The colorectum dissection image (left) and H&E results (right) of the healthy mice. K) Luminescence intensity of the CRCsensor from different organs of the healthy mice under 808- and 980- nm laser excitation.

A causal relationship between chronic colitis and an increased risk of CRC has been well-confirmed in numerous reports. However, the precise point at which inflammation gives way to carcinoma remains unclear, which significantly impedes fundamental research into the aetiology of the colorectum tumor and endangers the lives of patients in clinic. In order to elucidate the critical temporal point of colorectum carcinogenesis in vivo, the CRCsensor was employed to non-invasively detect miRNA-21 in the colorectum of mice with varying stages of chronic colitis (Figure 4A). The colitis-associated CRC mice model was established by the Azomethane (AOM)/Dextran sulphate sodium (DSS) chemical induction strategy in C57BL/6J mice for 10 weeks. The AOM is capable of inducing the formation of DNA O6-methylguanine adducts, resulting in a guanine (G) → adenine (A) transition, the primary mechanism by which genetic mutations are induced. The DSS has been demonstrated to impact DNA synthesis, impede epithelial cell proliferation, compromise the intestinal mucosal barrier, impair macrophage function and disrupt the intestinal flora, and may precipitate ulcerative colitis, which may in turn give rise to CRC when combined with the AOM.

**Figure 4.**
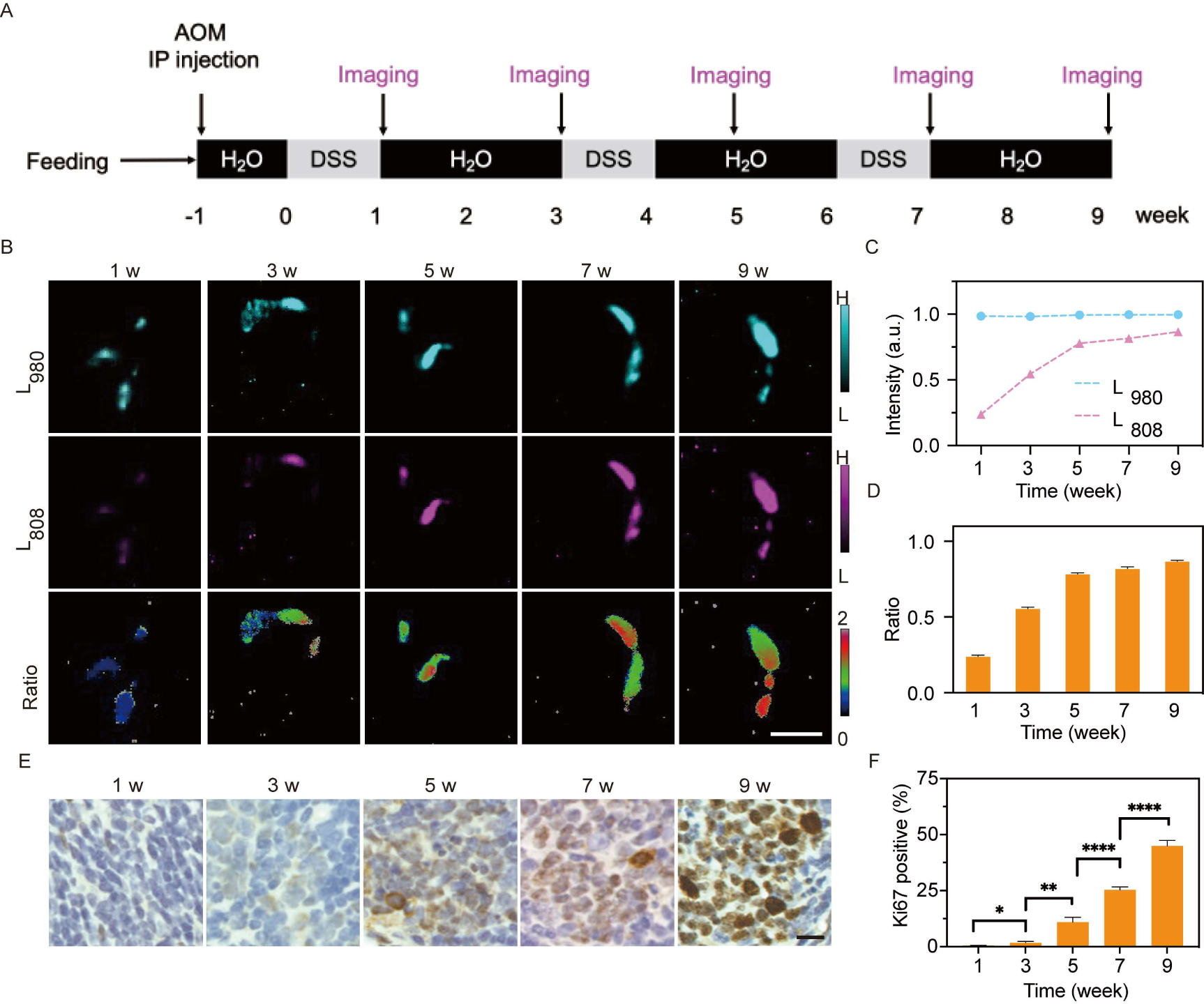
A) Schematic diagram of NIR-II ratio luminescence imaging of the mice with different stage of chronic colitis. B) Representative in vivo NIR-II luminescence and ratiometric images of mice with different stage of chronic colitis. Scale bars, 1.0 cm. C-D) NIR-II luminescence intensity of CRCsensor from the mice with different stage of chronic colitis. under 808- and 980- nm laser excitation (C) and the corresponding calculated ratio values (D). E) Immunohistochemical (IHC) results of the mice colorectum with different stage of chronic colitis and the corresponding calculated percentage of the Ki67 positive cells (below right). Scale bar: 10 μm. P values are indicated as follows: <0.05 (*), <0.01 (**), <0.001 (***), <0.0001 (****).

The CRCsensors were delivered to C57/BL6J mice with chronic colitis at varying stages, and the mice were imaged in the NIR-IIb region (Figure 4B). The responsive luminescence intensity observed under the 808 nm excitation channel demonstrated an increase in line with the duration of inflammation, while the luminescence intensity of the 980 nm excitation channel remained relatively consistent across different stages of chronic colitis in the mice model (Figure 4C, 4D). It is worth to note that the increasing rate of L808/L980 ratio in the first five-week period was markedly higher than that observed in the subsequent period, suggesting that the expression level of miRNA-21 in the early stages of chronic colitis cancerization was markedly elevated and then maintained at a high level. Due to the reduced scattering and deep penetration ability of CRCsensor NIR-IIb luminescence, the in vivo ratio luminescence was in good agreement with the in vitro results (Figure S24).

In vivo luminescence imaging can detect early cancerization in chronic colitis disease, as confirmed by ex vivo H&E sample analysis (Figure S25). No visible hyperplastic tissue was observed during the first week, indicating that no cancer had occurred. During the period of 1∼5 week, the H&E sections showed obvious hyperplastic tissue, but no obvious tumor tissue could be seen from the results of colorectal dissection, thus it is difficult to determine whether cancer has occurred. The obvious cancer tissues could be observed after week 7 from the dissection image and H&E analysis results of the mice colorectums, indicating that the inflammation has transformed into CRC. The colitis-associated cancerization process was further characterized by a classic in vitro immunohistochemical (IHC) analysis of colorectal tissue stained with Ki-67 protein, which has the ability to reflect the proliferative activity of cells in the tissue (Figure 4E). A gradual increase in proliferative activity was observed from the first week (0.44%) to the ninth week (44.9%), indicating a gradual transformation process from an inflammatory to a neoplastic state. Compared to week 5, the cell proliferative activity notably increased to 25.29% in week 7, which providing compelling evidence of tumor occurrence and was consistent with the colorectum dissection results. From the above results of the in vitro sample analysis, the occurrence of cancer could be clearly determined at the seventh week. The proposed strategy of luminescence imaging of miRNA-21 in the colorectum has been shown to detect early CRC at the third week of chronic colitis, representing a ∼ 4 weeks earlier detection than that achieved by the biopsy method through tissue sampling and the dissection observation through the naked eye.

## Conclusion

In summary, the risk of colorectal cancer can be effectively reduced by early and regular testing to detect cancerous changes in chronic colitis. We have developed a NIR-IIb luminescence nanosensor for imaging early CRC based on the monitoring of miRNA-21 expression levels in the colorectum. The CRCsensor can be split into two distinct entities, thereby allowing the IR820 dye to be released from the ErNPs, exhibiting a gradual recovery of NIR-II ratiometric luminescence due to the frustration of the ACIE process between ErNPs and the competing absorber fluorophore IR820. In the in situ CRC murine models, we demonstrated the capacity of the CRCsensor to facilitate precise non-invasive in vivo monitoring of CRC. This luminescence imaging technique enables the sensitive discrimination of early-stage CRC, which was achieved ∼ 4 weeks earlier than the clinical gold standard tactic (biopsy) in colitis-associated CRC models and observation by naked eye (endoscopy). Given that our strategy is non-invasive, it is clearly appropriate for periodic screening of early CRC, as well as for the evaluation of cancerization progression in chronic colitis. Furthermore, the administration of the CRCsensor via cystectomy enables a more rapid and secure arrival at the intended site in comparison to the intravenous or oral route. Due to the natural flexibility and programmability of DNA, our CRCsensor can be used to detect other disease markers (e.g. carcinoembryonic antigen) in the gut by replacing the target recognition sequence, which will greatly broaden the application of the sensor.

## Acknowledgements

This work was supported by the National Key R&D Program of China (2023YFB3507100), National Natural Science Foundation of China (22104017, 22088101, 22474026), New Cornerstone Science Foundation through the XPLORER PRIZE, and the Research Program of Science and Technology Commission of Shanghai Municipality (24QA2706400, 21142201000, 22JC1400400, 20490710600) and Innovation Program of Shanghai Municipal Education Commission (2023ZKZD08). Keywords: Colitis-associated colorectal cancer • MicroRNA • NIR-II luminescence • in vivo imaging • DNA nanosensor

## Table of Contents

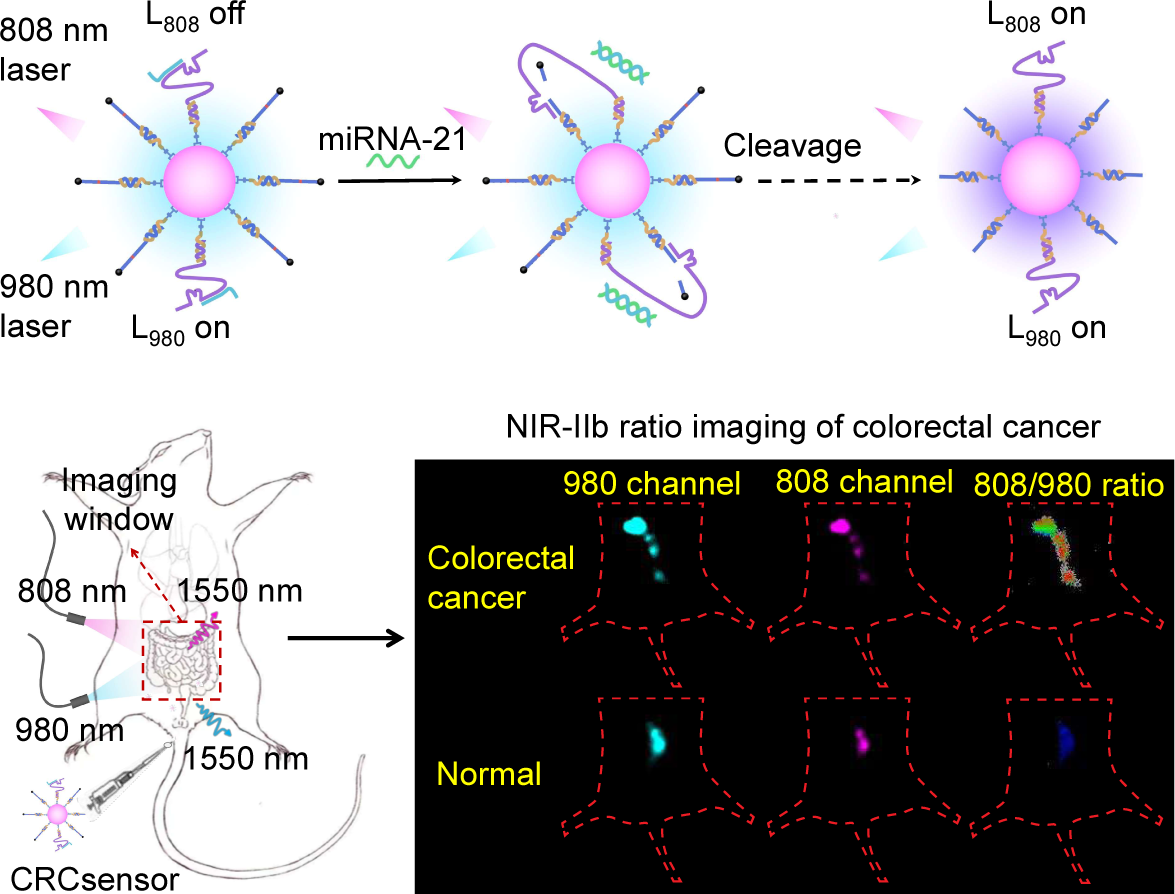

We reported a cascade amplified sensing strategy in the second near-infrared b (NIR-IIb, 1500-1700 nm) window for non-invasively in situ visualization of early cancerization biomarker miRNA-21. This strategy enabled a non-invasive detection of colitis-associated cancerization up to ∼4 weeks ahead of the clinically used biopsy analysis, providing a promising alternative for early diagnosis of chronic colitis cancerization and a guidance for therapeutic management.

