## Supporting Information for "Non-invasive diagnosis of early chronic colitis cancerization via amplified sensing of miRNA-21 in NIR-IIb window"

### Experimental procedures

#### Chemicals

Nucleic acids were obtained from Sangon biotech (Shanghai) Co., Ltd. Erbium (III) chloride hexahydrate, Yttrium (III) chloride hexahydrate, sodium trifluoroacetate (Na-TFA), 1-octadecene (ODE) and oleic acid (OA) were purchased from Sigma-Aldrich (St Louis, MO, USA). 1-ethyl-3-(3-dimethyl aminopropyl) carbodiimide hydrochloride (EDC•HCl) and N-hydroxy succinimide (NHS) were purchased from J&K Scientific Ltd. 4-mercaptobenzoic acid (MBA) and alendronic acid (ADA) were purchase from Aladdin Co., Ltd. (Shanghai, China). Dulbecco's modified eagle medium (DMEM) was purchased from MesGen Biotech Co., Ltd. (Shanghai, China). All chemicals were used as received without any further purification.

#### Characterizations

Transmission electron microscopy (TEM) tests were performed using a JEM-2100F transmission electron microscope with an accelerating voltage of 120 kV. Luminescence spectra were acquired on an Edinburgh FLS980 luminescence spectrometer with excitation of the 808- and 980- nm laser. Absorption spectra were collected by using a PerkinElmer Lambda 750S UV-visible-NIR spectrometer. The NIR-II in vivo luminescence imaging was performed with a modified home-built InGaAs array detector (Princeton Instruments, NIR vana 640). Dynamic light scatting tests were performed using a Malvern Zetasizer Nano ZS90 (Malvern Panalytical, UK)

#### Preparation of rare-earth trifluoroacetate precursors for the ErNP

5 mmol Rare-earth oxides were loaded into a flask containing 10 ml trifluoroacetic acid aqueous solution (v/v = 1:1). The slurry was heated to 70 °C and maintained until an optically transparent solution was formed. The obtained solution was filtered and condensed, and the resulting trifluoroacetate was dried at 100 °C for 12 h in a vacuum drying oven to obtain rare-earth trifluoroacetate powders.

#### Synthesis of hexagonal-phase $NaErF_4$ core nanoparticles.

The precursors (1.0 mmol $ErCl_3$) were added to a flask containing OA (17.00 mmol) and ODE (40.00 mmol). The mixture was heated to 140 °C under a vacuum until a clear solution was formed, after which the solution was cooled to 50 °C. Afterwards, 7.0 ml methanol solution containing ammoniumfluoride (4.0 mM) and sodium hydroxide (2.5 mM) were added, and the resultant solution was stirred for 60 min. After removing methanol, the solution was heated to 290 °C and maintained for 1.5 h under an argonatmosphere and then cooled to room temperature. The hexagonal-phase core nanoparticles were obtained by centrifugation at 5000 rpm for 5 min, washed with ethanol three times and re-dispersed in 10 ml of cyclohexane.

**Synthesis of hexagonal-phase core–shell-structured β-$NaErF_4$@$NaYF_4$ nanoparticles.**

A 2.5 ml amount of hexagonal core nanoparticle dispersion was added into a flask containing the shell precursors (0.5 mmol Na-TFA and 0.5 mmol Y-TFA) and solvents (20.0 mmol OA and 20.0 mmol ODE). The slurry was heated to 100 °C under a vacuum for 30 min to remove cyclohexane, water and oxygen, followed by heatingto 290 °C for 90 min under an argon atmosphere. The after treatments are the same as those to get the $NaErF_4$ core nanoparticles.

**Synthesis of surface modified water soluble ErNP@ADA-MBA@linker DNA**

A ligand exchange reaction was conducted to functionalise ErNPs with an amine group and disperse them in an aqueous solution. A solution of 0.5 mL of the aforementioned dispersed RENPs (100 mM) in cyclohexane was prepared by adding 5 mg of $NOBF_4$ to a centrifuge tube and shaking for 2 minutes. Following a five-minute period of rest, the supernatant cyclohexane was discarded, and 3 mL of isopropanol was added. The mixture was subjected to centrifugation at 15,000 rpm for 10 minutes and subsequently dissolved in 0.1 mL of ultrapure water. The water-soluble ErNPs were added to a 5 mL solution of $H_2O$/water containing 10.00 mM ADA and 10.00 mM MBA, and the mixture was stirred overnight. The ErNP@ADA-MBA was then washed with ultrapure water on three occasions and re-dispersed in 10 mL of ultrapure water.
1.0 mL of the ErNP@ADA-MBA was mixed with 50 uL 100 uM linker DNA solution and shaken overnight. The as-obtained ErNP@ADA-MBA@linker DNA was centrifuged, washed with Tris-HCl buffer (10 mM, containing 100 mM KCl and 1.0 mM EDTA, pH 8.0) for three times and re-dispersed in 1 mL Tris-HCl buffer for future use.
The number of ErNP@ADA-MBA@linker DNA ($N_{Er}$) in the solution was counted by nanoparticle tracking analysis (NTA) on a *Malvern NanoSight NS300* system, while the molar concentration of the substrate and DNAzyme were determined by measuring luminescence spectrum of 6-Carboxyfluorescein (6-FAM) modified substrate and DNAzyme.

**Preparation of IR820 labelled DNA substrate strand (Sub-IR820)**

50 μM the thiol-modified DNA substrate strand (100 μM) was mixed with 1.0 mL Tris(2-carboxyethyl)phosphine (TCEP, 1 mM) and shaken for 1 hour. Thereafter, the obtained solution was mixed with 1mL 5 μM IR820 DMF solution and shaked for 12

hours.The as-obtained Sub-IR820 was precipitated with 75% ethanol (10 mL) and 1M NaCl (1.0 mL) in -20 °C for 4 h. The mixture was subjected to centrifugation at 15,000 rpm for 10 minutes and subsequently dissolved in 0.1 mL of Tris-HCl buffer.

**Synthesis of the CRCsensor**

Equal amount of the DNAzyme strand and the locker strand was mixed in Tris-HCl buffer and then heated to 95 °C for 5 min, followed by slow cooling to room temperature to obtain DNAzyme&locker. 1.0 mL of the ErNP@ADA-MBA@linker DNA was mixed with 100 uL Sub-IR820 (100 μM) and 100 μL DNAzyme&locker (25 μM) solution and shaken at 30 °C for 6 hours. The as-obtained CRCsensor was centrifuged re-dispersed in 1 mL Tris-HCl buffer for future use.

**Polyacrylamide gel electrophoresis (PAGE) analysis**

12% polyacrylamide gel was prepared by mixing 3.0 mL water, 2.4 mL 5xTBE buffer, 6 mL 30 % acrylamide, 50 μL APS and 25 μL TEMED. 10 uL 1.00 μM DNA sample and 1.5 μL 6x DNA gel loading buffer were mixed and injected into the lane. The voltage of gel electrophoresis was set as 120V. After running 90 min in 1xTBE buffer, the polyacrylamide gel was stained with 50 mL 1xTBE buffer containing 15 μL 4S gel red for 30 min. The polyacrylamide gel was scanned with a luminescence gel imaging system.

**In vitro characterization of CRCsensor respond to miRNA-21**

The CRCsensors were incubated with miRNA-21 in Tris-HCl buffer at 37 °C for 4 h. Luminescence of the samples was monitored by luminescence spectrometer and CCD with 808- and 980- nm lasers.

**Cell culture**

MC38 cells and MCEC cells were cultured in DMEM containing 10% FBS, 80 U/mL penicillin and 0.08 mg/mL streptomycin. All cells were maintained at 37 °C in a humidified incubator containing 5% $CO_2$ and 95% air.
To simulate intracellular miRNA-21 expression difference, a DNA strand with a complimentary sequence matching that of miRNA-21 (miRNA-21 inhibitor) and a DNA strand with a sequence analogous to that of miRNA-21 (miRNA-21 mimics) were pre-transfected into MC38 cells and MCEC cells via lipofection 3000, respectively.

**Cytotoxicity test**

The cytotoxicity of CRCsensor was evaluated using the Cell Counting Kit-8 (CCK8) assay. Once the cell density reached 80-90%, the cells were digested, centrifuged and dispersed into 20 mL DMEM. 200 μL of the obtained cell suspension was seeded in a 96-well plate and maintained at 37°C overnight. Subsequently, the cell culture DMEM was replaced with 140 μL DMEM containing various concentrations of CRCsensor, and the cells were incubated for 12 h (five parallel wells were set for each CRCsensor concentration). 50 μL of CCK8 solution was added to each well and the cells were incubated for 4 h. The optical density (OD) was then measured at 450 nm via microplate reader.

**Total miRNA extraction**

Total miRNAs were extracted from MC38 and MCEC cells by the SanPrep Column microRNA Extraction Kit according to reagent protocol. In brief, 0.2 ml of chloroform was mixed with the cell lysate and centrifuged at 12,000 rpm for 10 min. The upper aqueous phase was transferred to a new tube and 1/3 volume of anhydrous ethanol was added. The solution was added to an adsorbent column and centrifuged at 12,000 rpm for 2 min. The obtained flow-through solution was subsequently collected and mixed with 2/3 volume of anhydrous ethanol, following by centrifuging at 12,000 rpm for 2 min. The column was rinsed with RPE solution and centrifuged at 10,000 rpm for 30 s. A volume of 30 μL of RNase-free water was added to the adsorbent membrane and centrifuged at 12,000 rpm for 2 min to collect the miRNA solution. The obtained miRNA solution was stored at -70°C for further use.

**In vitro luminescence imaging of CRCsensor in cells**

NIR-II luminescence microscopy imaging was performed via a home-build NIR-II microscope with 808- and 980- nm excitation and 1500-1700 nm as detection window. 150 μL opti-MEM containing 10 μL CRCsensor were incubated with cells for 4 h, washed with PBS for two times, The cells were immersed with 4% paraformaldehyde (PFA) fix solution and imaged under the NIR-II luminescence microscope.

**Establishment of mice subcutaneous CRC model**

25 μL matrix gel was thawed and combined with 25 μL MC38 cell suspension. The resulting mixture was then injected subcutaneously into the hind limbs of C57BL/6J mice after dehairing.

**In vivo luminescence imaging of the CRCsensor in subcutaneous CRC mice models**

The subcutaneous tumour-bearing C57BL/6J mice were anaesthetised and the hair was shaved prior to luminescence imaging. Subsequently, CRCsensor (0.1 mM in saline based on $Er^{3+}$, 50 μL) was in situ injected into the tumour. Thereafter, in vivo NIR-II luminescence images (1500 LP) of the mice were acquired over time using an InGaAs CCD camera, with excitation by 808- and 980-nm lasers, respectively.

**Establishment of mice in situ colorectal cancer model**

An in situ colorectal cancer model was established using the chemoinduction method. C57BL/6J male mice (12 weeks old) were intraperitoneally injected with azoxymethane (AOM) at a dose of 10 mg/kg, and subsequently alternated between dextran sulfate (DSS) solution (2%) and normal drinking water. The DSS treatment cycle was typically one week, followed by 2 weeks of normal drinking water, and this cycle could be repeated 3 times.

**In vivo luminescence imaging of the sensor in orthotopic CRC mice models**

The orthotopic tumour-bearing C57BL/6J mice were anaesthetised and the hair was shaved prior to luminescence imaging. Subsequently, CRCsensor (0.1 mM in saline based on $Er^{3+}$, 50 μL) was administered into the colorectum of the mice via pipette. Thereafter, in vivo NIR-II luminescence images (1500 LP) of the mice were obtained using an InGaAs CCD camera over time with excitation by 808- and 980-nm lasers, respectively.

**Sample preparation for pathologic and immunohistochemical analysis**

For pathological analysis, the colons were excised and immersed in PFA for 24 hours. The sample was then dehydrated in an ethanol solution, embedded in paraffin, and cut into sections. The paraffin was removed by xylene washing, and the sections were incubated with haematoxylin for four minutes and eosin for two minutes (H&E), followed by washing with distilled water.

For immunohistochemical analysis, the samples were subjected to a dehydration process and subsequently embedded in paraffin wax, after which they were cut into sections. Subsequently, the paraffin sections of the grafts were deparaffinised and placed in citrate buffer, which was then boiled in order to facilitate antigen retrieval. Subsequently, the sections were incubated in a 3% hydrogen peroxide solution at room temperature to block endogenous peroxidase activity. Subsequently, bovine serum albumin (BSA, 3%) was added dropwise to the section and incubated at room temperature for 30 minutes. The primary antibodies (Ki67) were added to the sample and incubated for eight hours at 4 °C. Following this, the sample was washed with PBS three times. Subsequently, the horseradish peroxidase (HRP) labelled secondary antibody was applied and incubated at room temperature for 30 minutes. Thereafter, the samples were subjected to DAB and haematoxylin staining for colouration

### Results and discussion

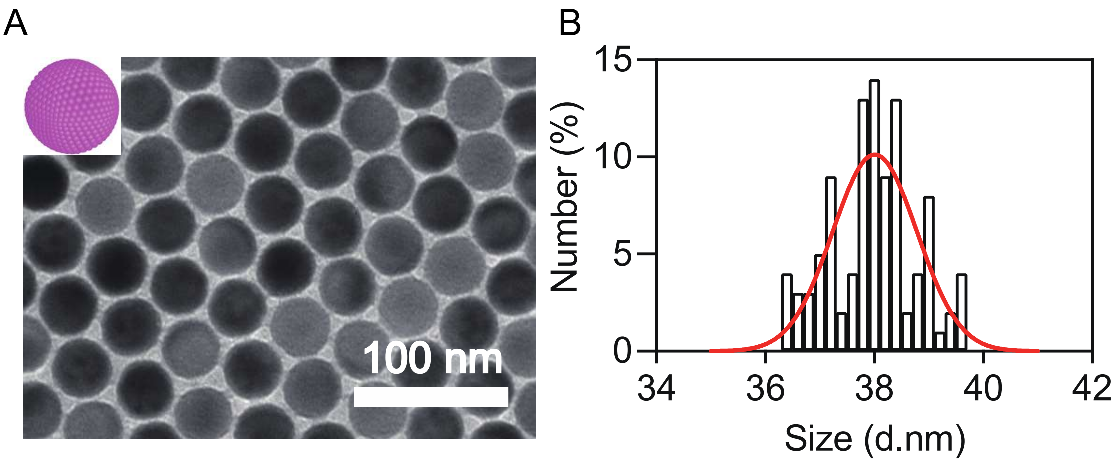

**Figure S1.** (A) Transmission electron microscopic image and (B) the size distribution of the $NaErF_4$ core.

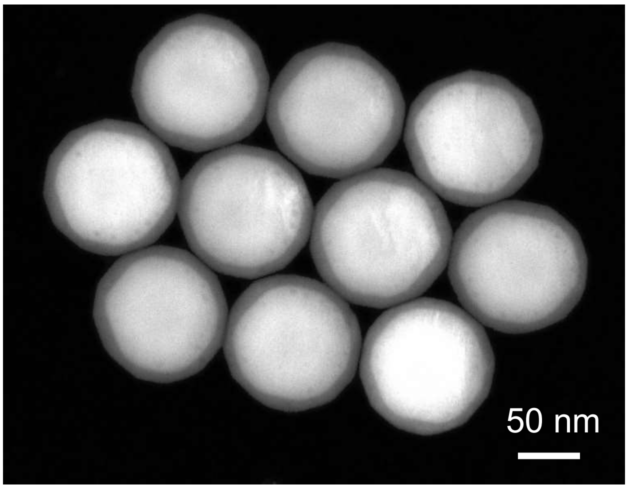

**Figure S2.** High angle annular dark field image of the $NaErF_4$@$NaYF_4$ core-shell particles.

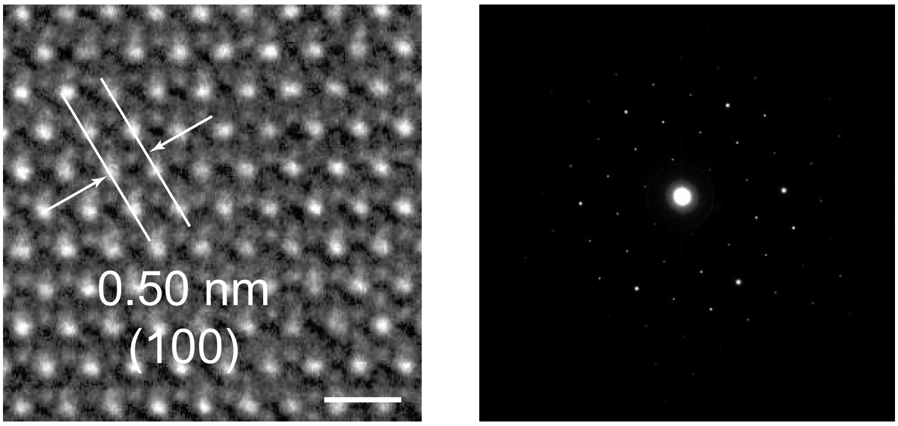

**Figure S3.** The high resolution TEM image of a single ErNP nanoparticle (left, scale bar: 1 nm) and the corresponding electron diffraction pattern (right).

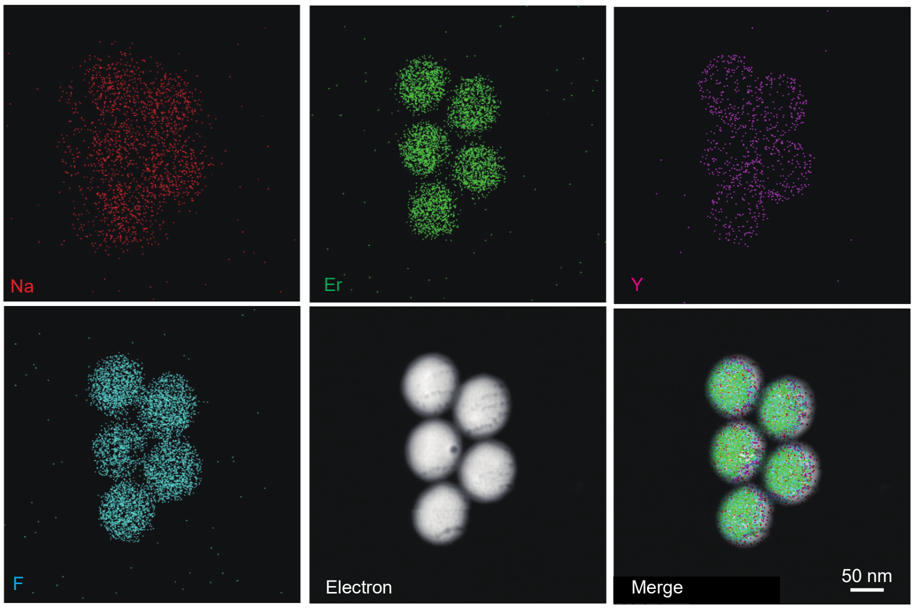

**Figure S4.** Elemental mapping images of the $NaErF_4$@$NaYF_4$ core-shell particles.

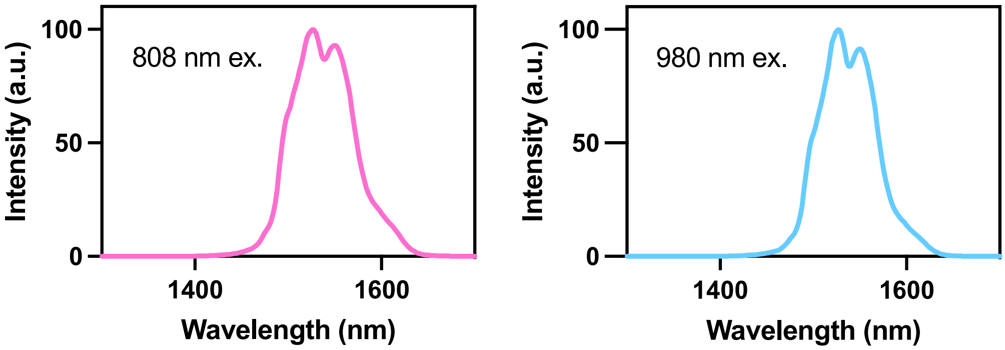

**Figure S5.** Luminescence emission spectra of $NaErF_4@NaYF_4$ under 808- (left) and 980- (right) nm laser excitation in cyclohexane phase.

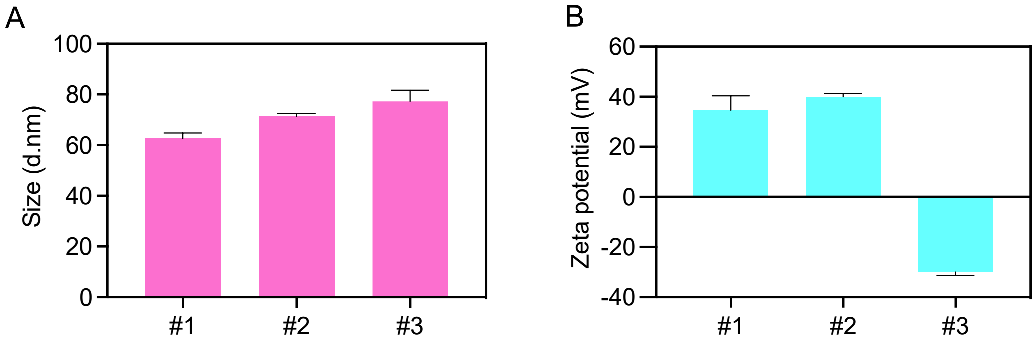

**Figure S6.** Hydrodynamic size (A) and zeta potential (B) of the $ErNP@BF_4^-$ (#1), the ErNP@ADA-MBA (#2) and the ErNP@ADA-MBA@linker DNA (#3).

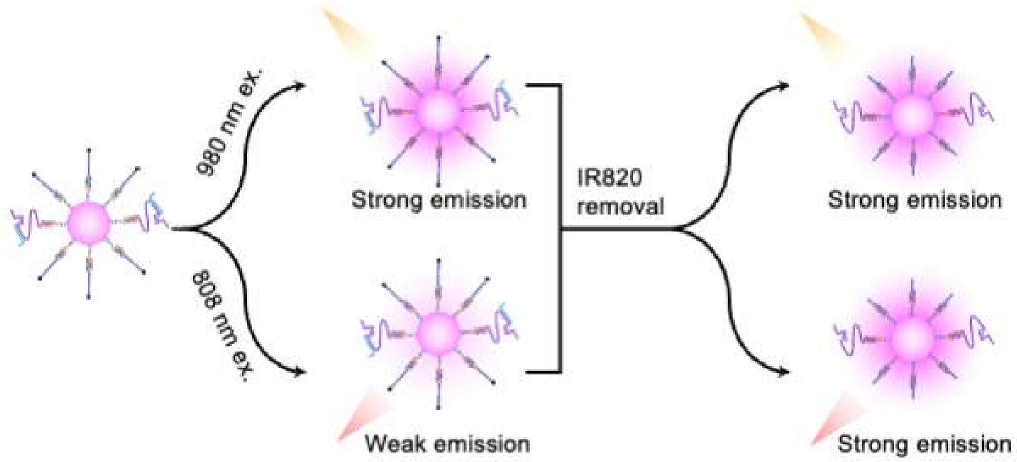

**Figure S7.** Schematic depiction of the the absorption competition-induced emission (ACIE) principle in this study.

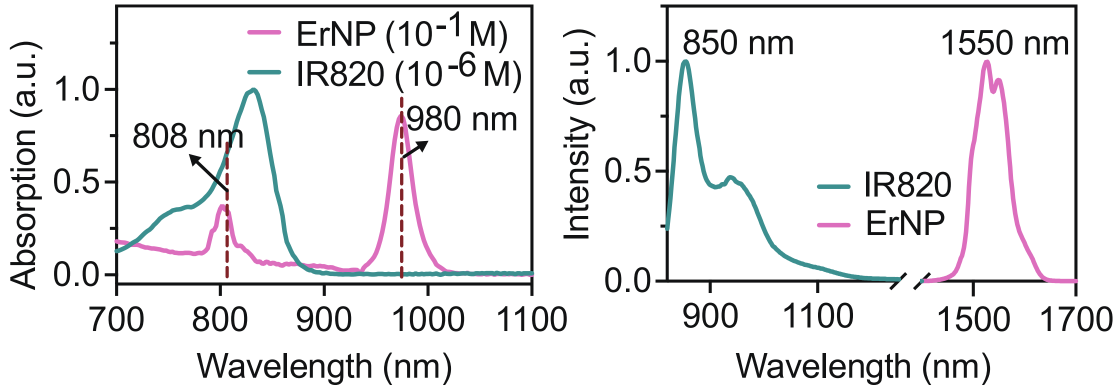

**Figure S8.** Luminescence emission spectra of ErNP and ErNP@IR820 under 980- (left) and 808- (right) nm laser excitation in DMF. Absorption spectra (left) and luninescence emission spectra (right) of the ErNP and the IR820 fluorophore. The concentration of ErNP was caluculated as $Er^{3+}$.

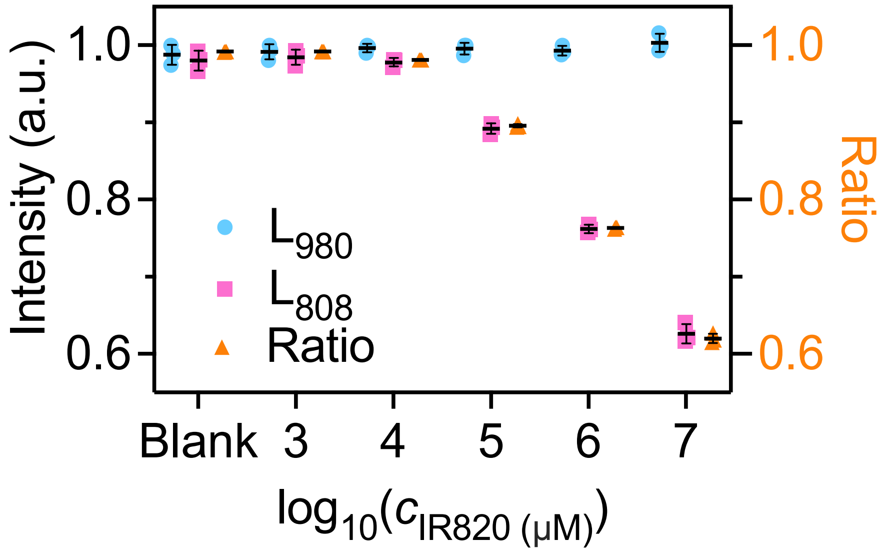

**Figure S9.** NIR-II luminescence intensities of ErNP (1 μM) coated with different concentration of IR820 dye under 808- and 980- nm laser excitation and the corresponding calculated ratio values.

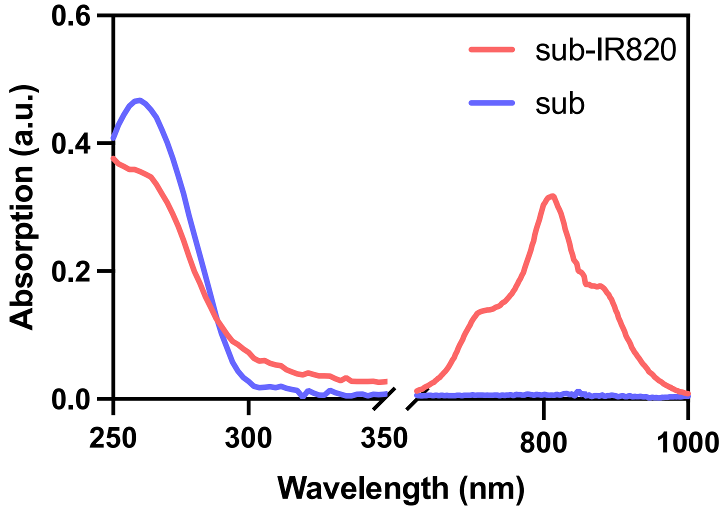

**Figure S10.** Absorption spectra of the substrate (sub) and the IR820 dye modified substrate (sub-IR820).

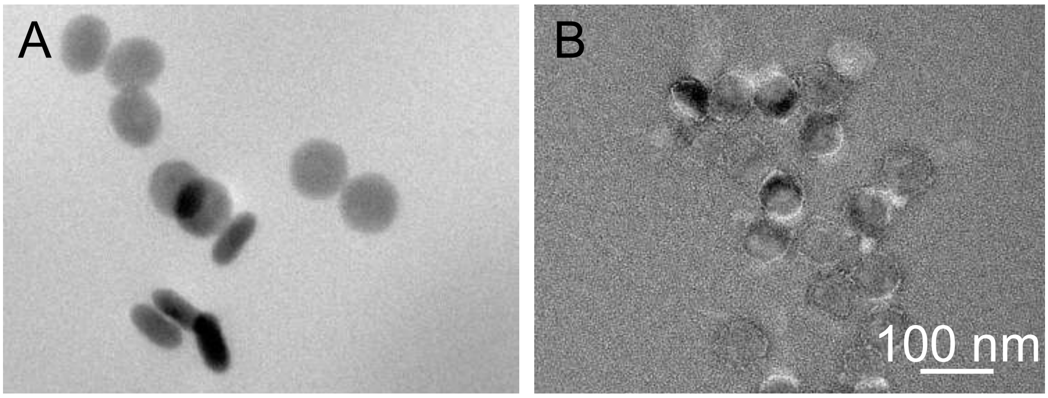

**Figure S11.** Transmission electron microscopic image of the CRCsensor (A) and the CRCsensor stained with 2% uranium acetate (B).

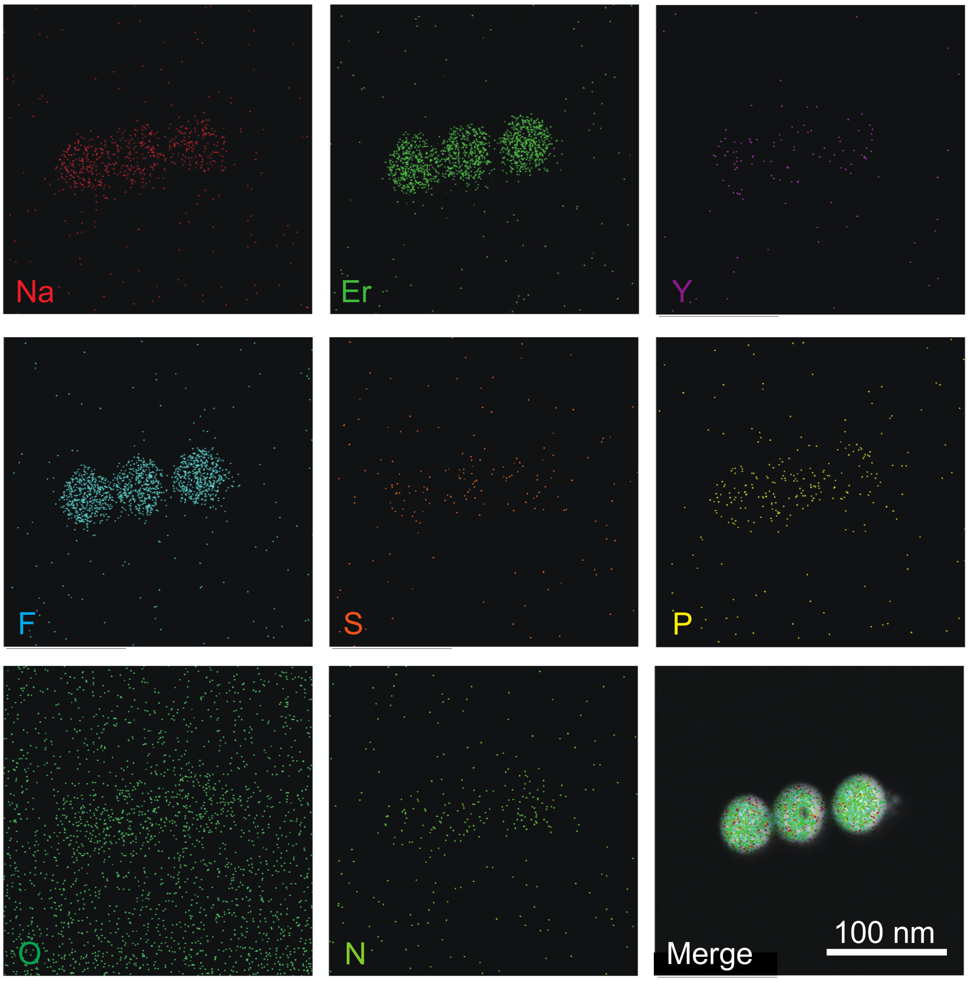

**Figure S12.** Elemental mapping images of the CRCsensor.

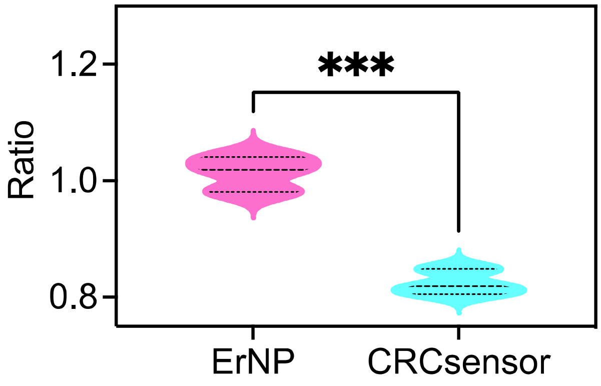

**Figure S13.** $L_{808}/L_{980}$ ratio of the ErNP and the CRCsensor.

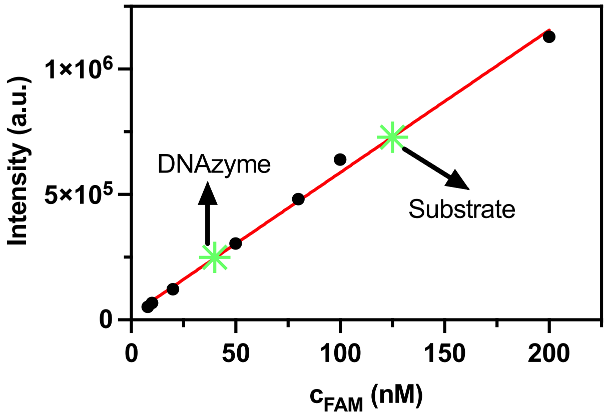

**Figure S14.** Quantification of substrate and DNAzyme attached on each nanoparticle.

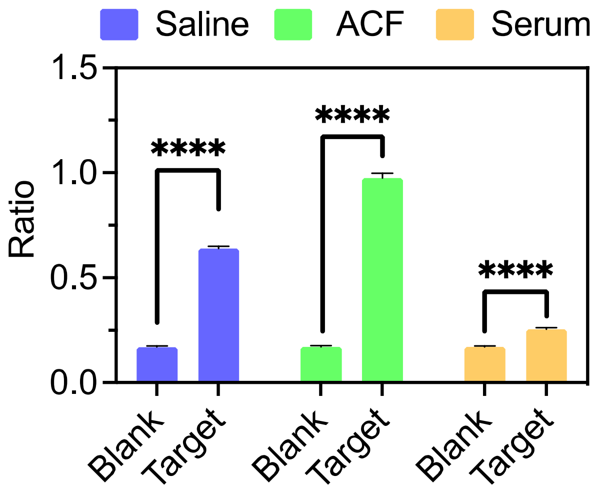

**Figure S15.** miRNA-21 responsive ability of the CRCsensor in complex biology samples. ACF: artificial colon fluid. P values are indicated as follows: <0.0001 (****).

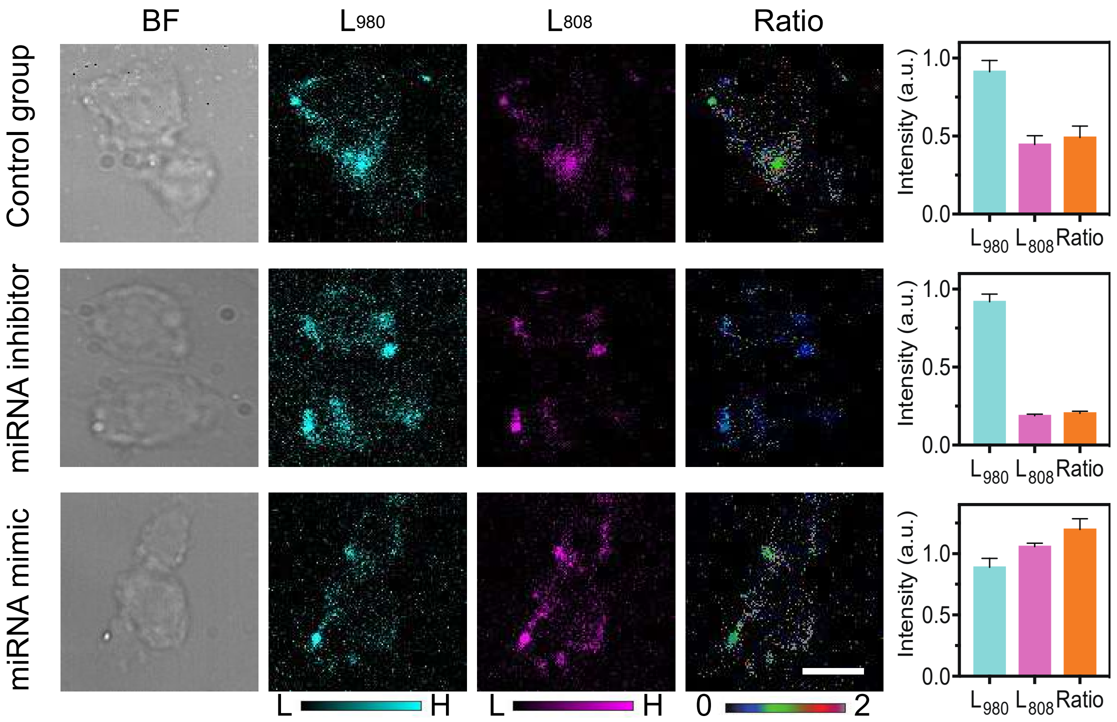

**Figure S16.** NIR-II luminescence imaging and the corresponding intensitis values of the CRCsensor incubated with MC38, MC38 pretreated with miRNA-21 inhibitor and MC38 pretreated with miRNA-21 minic (Scale bar: 25 μm).

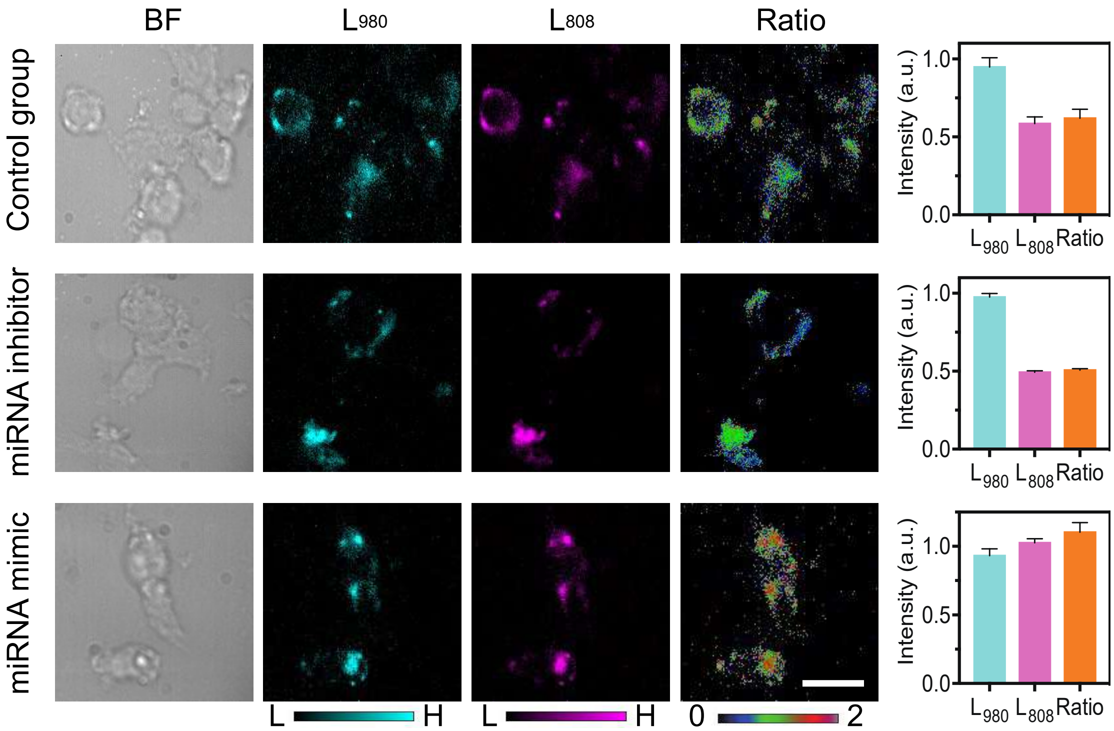

**Figure S17.** NIR-II luminescence imaging and the corresponding intensitis values of the CRCsensor incubated with MCEC, MCEC pretreated with miRNA-21 inhibitor and MCEC pretreated with miRNA-21 minic (Scale bar: 25 μm).

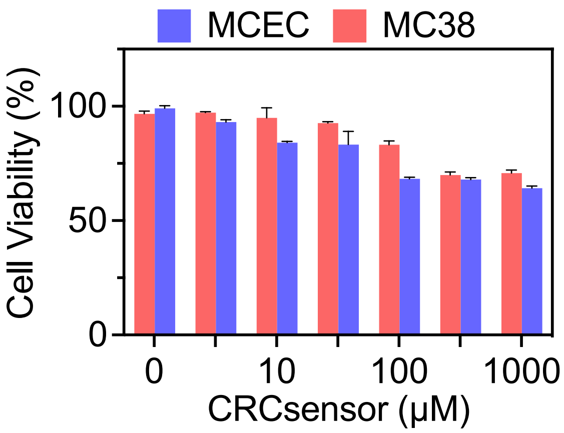

**Figure S18.** CCK8 assays of MCEC and MC38 cells treated with different concentrations of the CRCsensor (5-1000 μM).

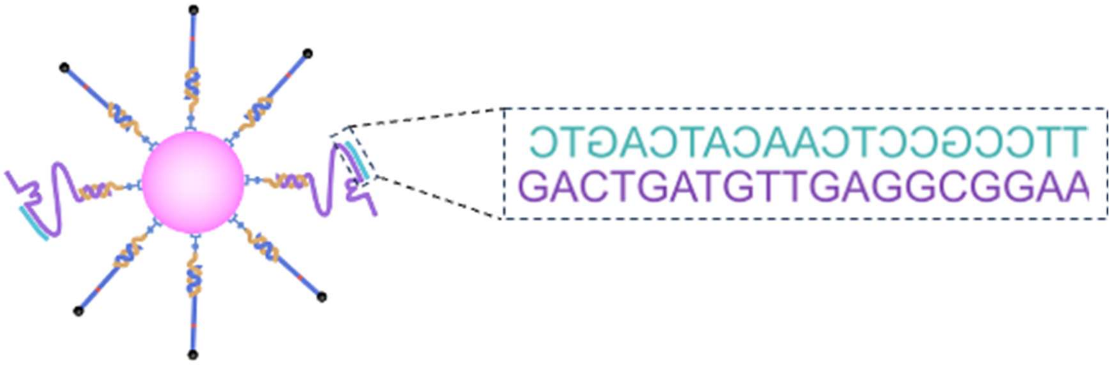

**Figure S19.** Scheme illustration and DNA sequence information of the non-responsive CRCsensor.

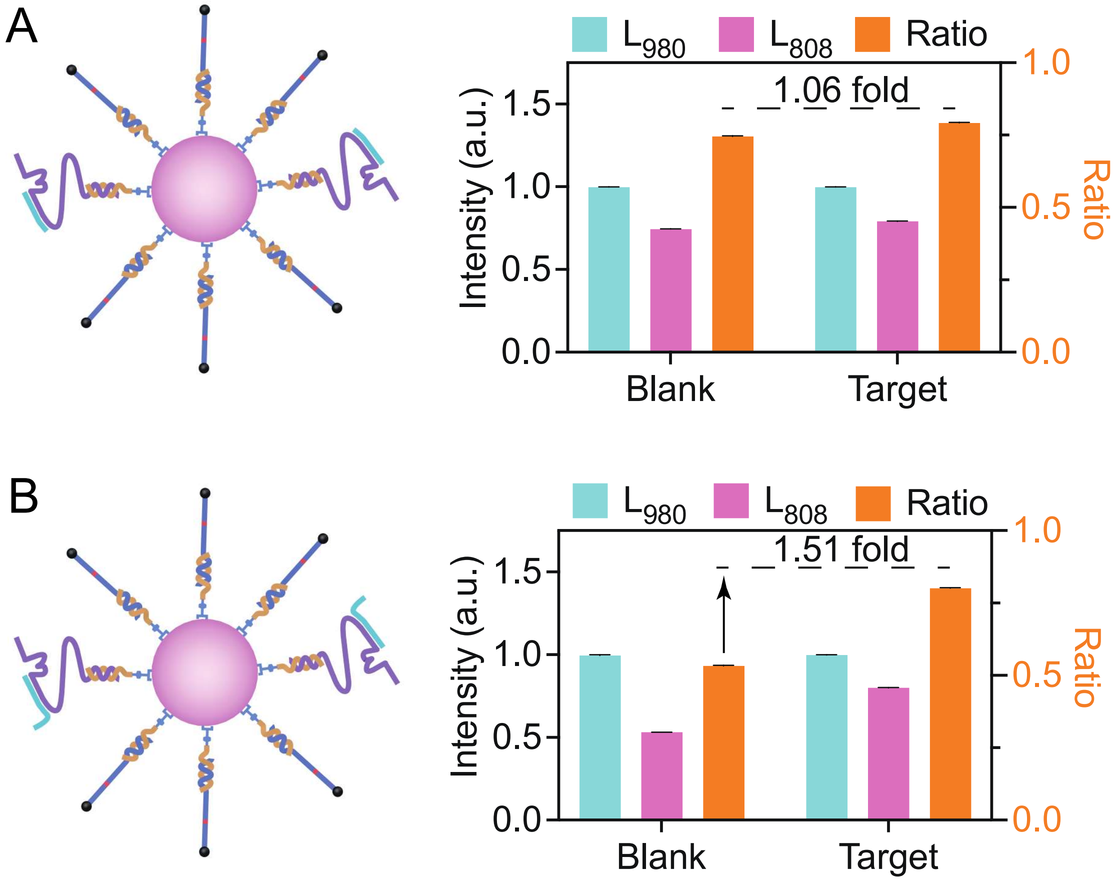

**Figure S20.** miRNA-21 responsive ability of the non-responsive CRCsensor (A) and the CRCsensor (B).

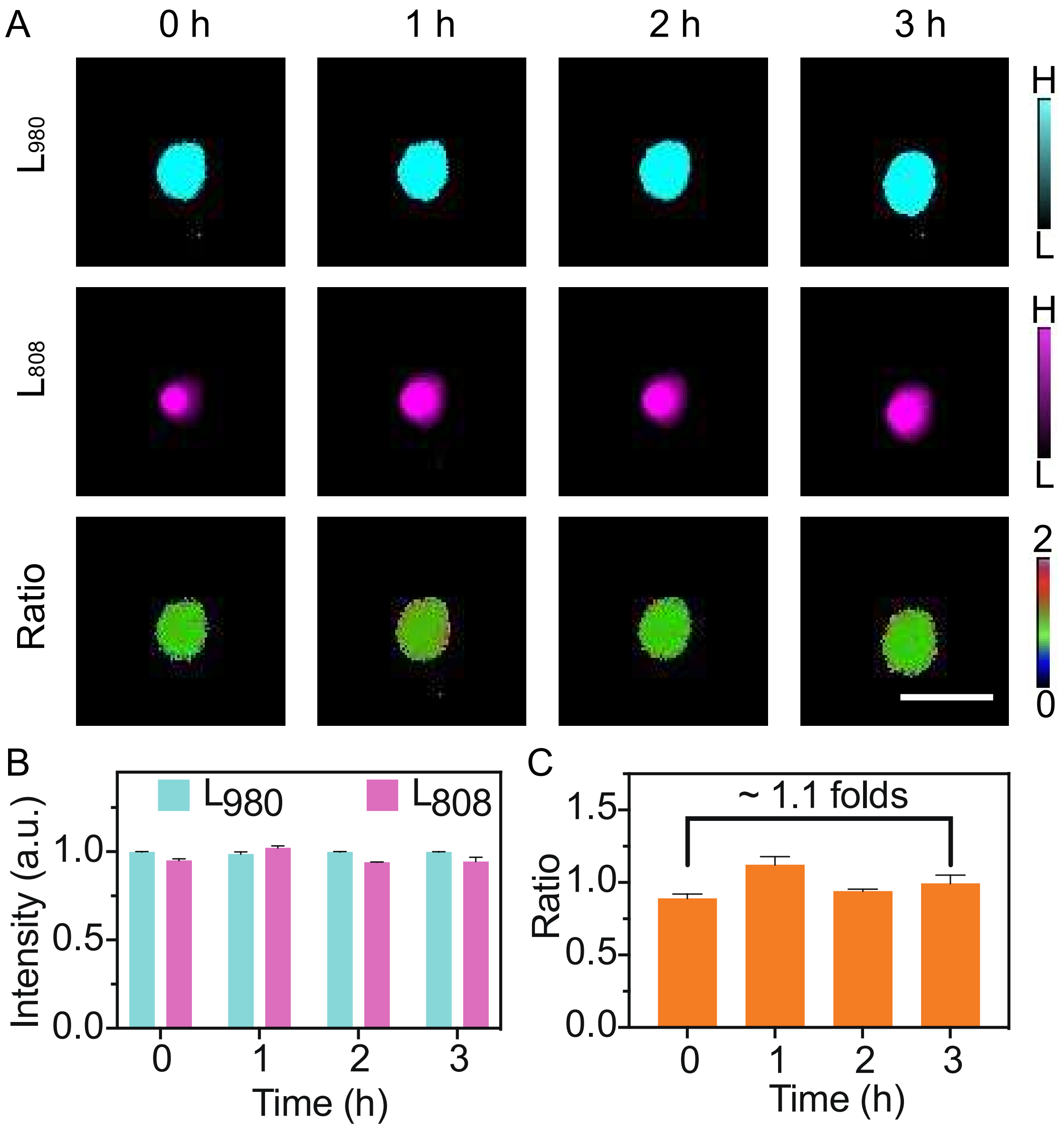

**Figure S21.** (A) Respresentive NIR-II luminescence imaging, (B) the corresponding intensitis values and (C) the calculated $L_{808}/L_{980}$ ratio of the non-responsive CRCsensor in the subcutaneous CRC-bearing mice. Scale bar: 1.0 cm.

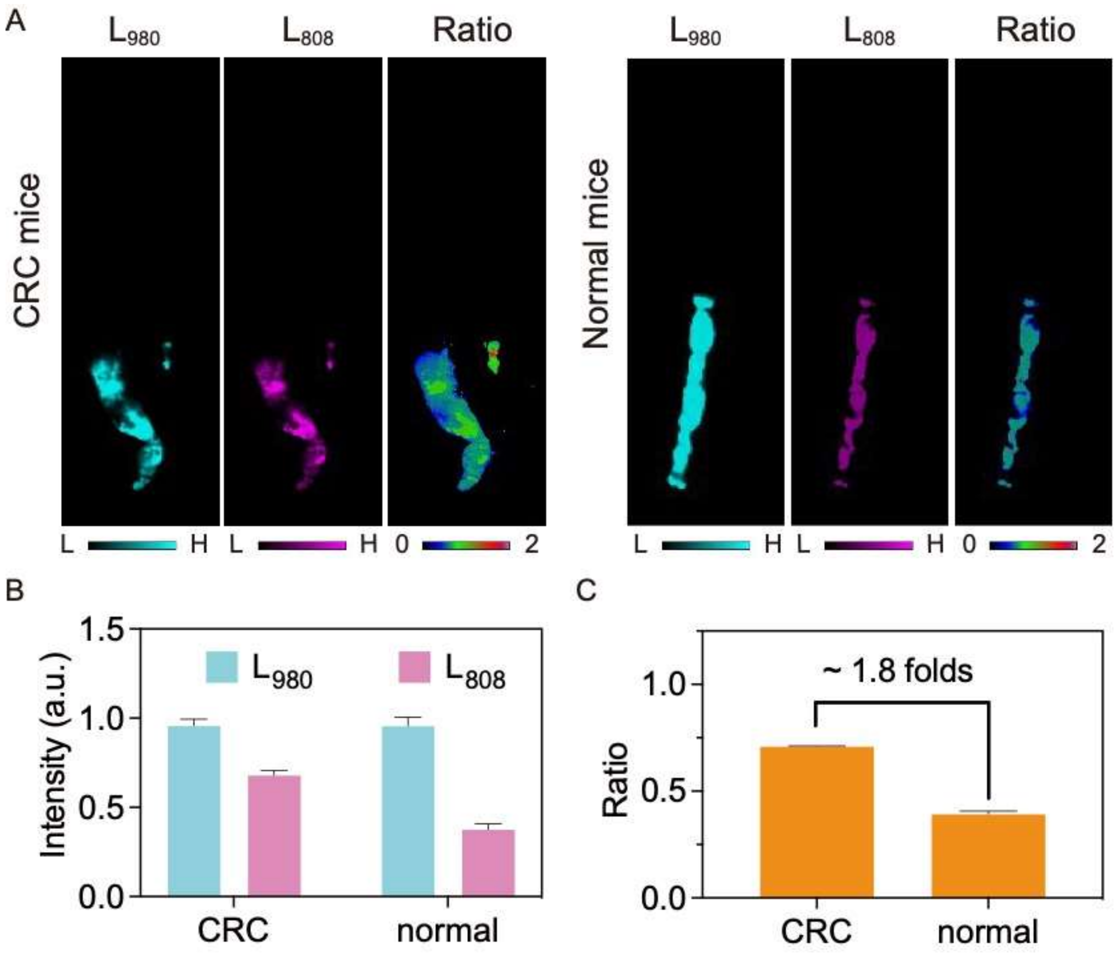

**Figure S22.** (A) Respresentive NIR-II luminescence and ratio images, (B) the corresponding intensitis values and (C) the calculated $L_{808}/L_{980}$ ratio of the colorectum tissues removed from the CRC-bearing mice and the normal mice.

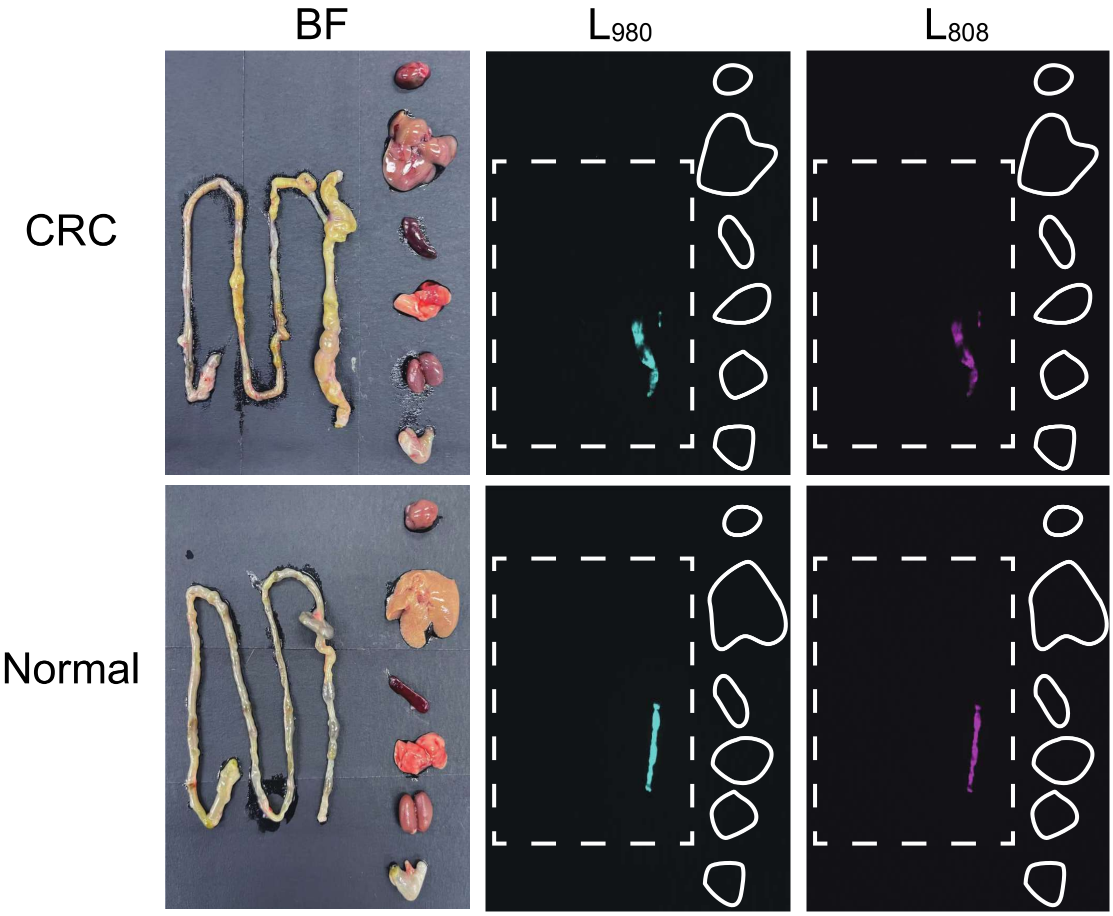

**Figure S23.** The dissection images and the corresponding NIR-II imaging results of CRCsensors distributions in different organs of CRC mice and normal mice. Left: intestinal; Right: heart, liver, spleen, lung, kidney, stomach (from top to down).

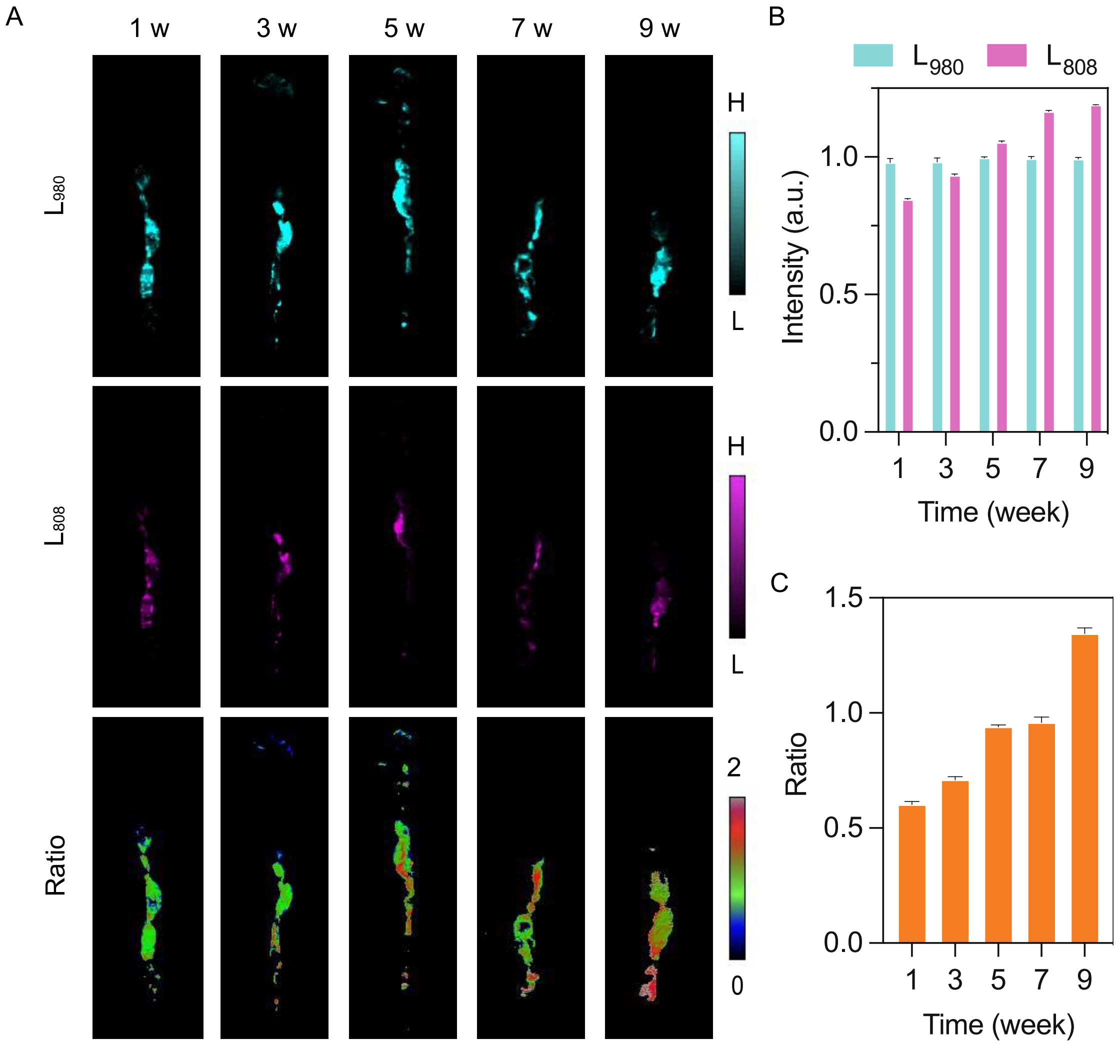

**Figure S24.** (A) NIR-II luminescence and ratio images of the colorectum, (B) the corresponding intensitis values and (C) the calculated $L_{808}/L_{980}$ ratio of mice with different stage of chronic colitis.

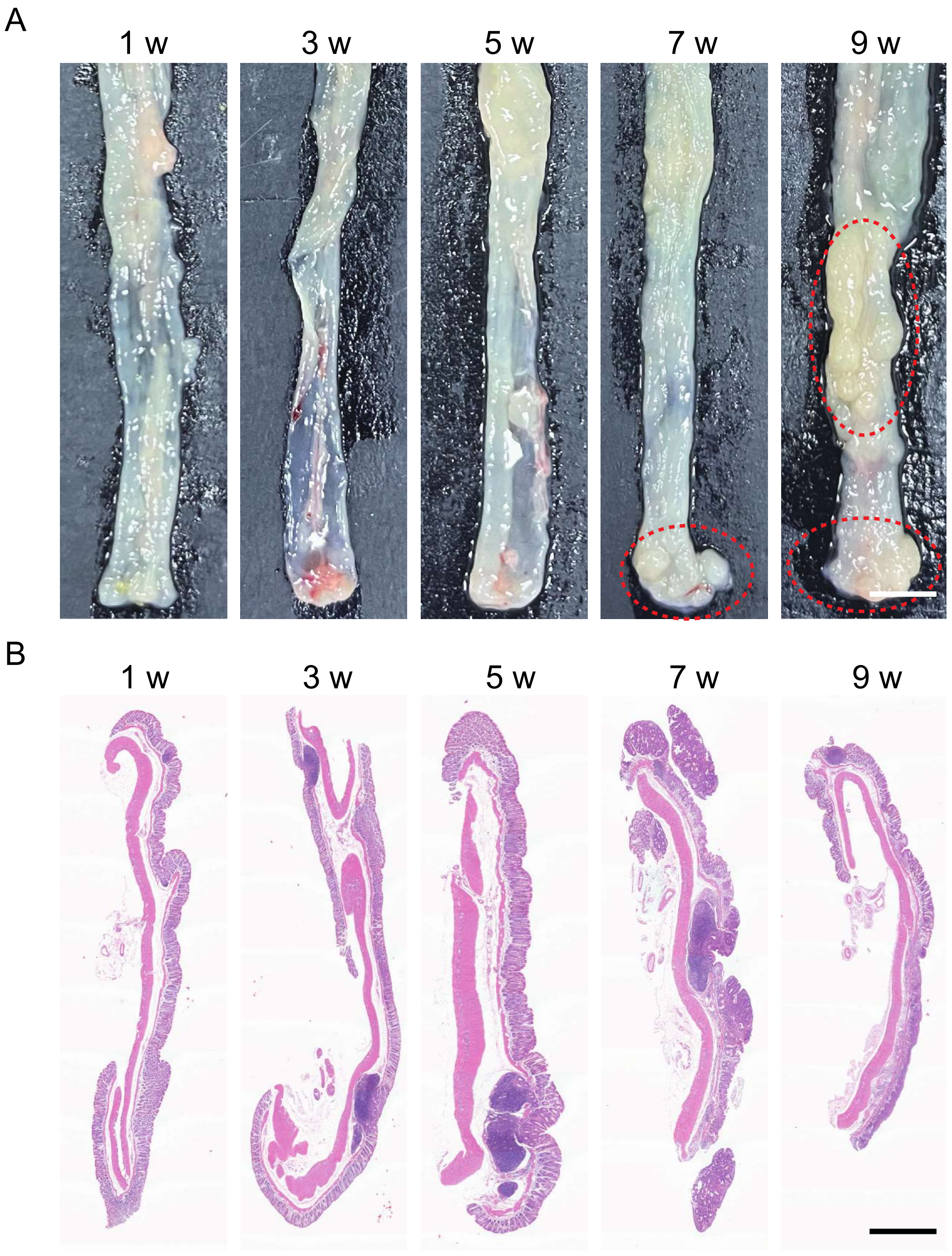

**Figure S25.** (A) The colorectum dissection images (scale bar: 5 mm, red circles: tumors) and (B) H&E results (scale bar: 1 mm) of mice colorectum with different stage of chronic colitis.

**Table S1.** DNA and RNA sequences used in this study.

| Oligonucleotide | Sequence (5’ to 3’) |
|---|---|
| Linker DNA | GGCTGGCTGGCTGGCTGGCTTTTTTT-maleimide |
| Sub-SHSH | AGCCAGCCAGCCAGCCAGCCTTTTTTTTTTTTCGATCTCCTATGG/rA/AAGTTCCGCCT-SHSH |
| DNAzyme | AGCCAGCCAGCCAGCCAGCCTTTTTTTTTTTTTTTTTTTTTTTTTTTTTTTGACTGATGTTGAGGCGGAA*CCAGGTCAAAGGTGGGTGAGGGGACGCCAAGAGTCCCCGCGGT*TAGGAGATCG |
| Locker | TTCCGCCTCAACATCAGTCTGATAAGCTA |
| Locker* | TTCCGCCTCAACATCAGTC |
| miRNA-21 | UAGCUUAUCAGACUGAUGUUGA |
| M1 | TAGATTATCAGACTGATGTTGA |
| M2 | TAGATTATCAGACTGACGTTGA |
| M3 | TAGCATATCAGTCTGATCTTGA |
| miRNA-155 | UUAAUGCUAAUCGUGAUAGGGGUU |
| miRNA-375 | UUUGUUCGUUCGGCUCGCGUGA |
| miRNA-21 mimic | TAGCTTATCAGACTGATGTTGA |
| miRNA-21 inhibitor | TCAACATCAGTCTGATAAGCTA |
